# Plasmalogen oxidation induces the generation of excited molecules and electrophilic lipid species

**DOI:** 10.1101/2024.02.22.581635

**Authors:** Rodrigo L. Faria, Fernanda M. Prado, Helena C. Junqueira, Karen C. Fabiano, Larissa R. Diniz, Mauricio S. Baptista, Paolo Di Mascio, Sayuri Miyamoto

**Author notes:** To whom correspondence may be addressed: Sayuri Miyamoto: **Email:**.

## Abstract

Plasmalogens are glycerophospholipids with a vinyl-ether linkage at the sn-1 position of the glycerol backbone. Despite being suggested as antioxidants due to the high reactivity of their vinyl ether groups with reactive oxygen species (ROS), our study reveals the generation reactive oxygen and electrophilic lipid species from oxidized plasmalogen intermediates. By conducting a comprehensive analysis of the oxidation products by liquid chromatography coupled to high-resolution mass spectrometry (LC-MS) we demonstrate that singlet molecular oxygen [O_2_ (^1^Δ_g_)] reacts with the vinyl ether bond, producing hydroperoxyl acetal as major primary product (97%) together with minor quantities of dioxetane (3%). Furthermore, we show that these primary oxidized intermediates lead to the formation of excited triplet carbonyls, O_2_ (^1^Δ_g_), and electrophilic phospholipid and fatty aldehyde species, as secondary reactive products. The generation of excited triplet carbonyls from dioxetane thermal decomposition was confirmed by light emission measurements in the visible region using dibromoantracene as a triplet enhancer. Moreover, O_2_ (^1^Δ_g_) generation from dioxetane and hydroperoxyacetal was evidenced by detection of near-infrared light emission at 1270 nm and chemical trapping experiments. Additionally, we have thoroughly characterized alpha-beta unsaturated phopspholipid and fatty aldehydes by LC-MS analysis using two probes that specifically reacts with aldehydes and alpha-beta unsaturated carbonyls. Overall, our findings demonstrate the generation of excited molecules and electrophilic lipid species from oxidized plasmalogen species unveiling the potential prooxidant nature of plasmalogen oxidized products.

**Significance Statement:** Plasmalogens, the most abundant subclass of ether lipids in mammalian cells, have traditionally been regarded as antioxidants. However, our study reveals a new perspective, shedding light on the generation of chemiexcited and reactive lipid species during plasmalogen photooxidation. We provide direct evidence revealing the production of excited triplet carbonyls and singlet molecular oxygen as secondary reactive products originating from dioxetane and hydroperoxyacetal intermediates. Importantly, we also demonstrate the generation of electrophilic alpha-beta unsaturated phospholipids and fatty aldehydes through plasmalogen oxidation. These findings highlight the production of excited states and reactive lipid species resulting from plasmalogen oxidation, which can potentially induce oxidative modifications in biological systems.

## Introduction

Plasmalogens are unique type of glycerophospholipids containing a vinyl-ether bond at the sn-1 position of the glycerol backbone and a sn-2 acyl chain enriched in polyunsaturated fatty acids (PUFA), such as arachidonic acid. Plasmalogens are the most abundant ether lipids found in all human tissues, especially in the brain and heart (1-3). They are also found in anaerobic bacteria, while absent in plants, fungi, and most aerobic bacteria (2). Constituting approximately 15-20% of the total phospholipid pool in mammals, plasmalogens serves as major structural membrane components, being important for membrane integrity and as a reservoir for lipid mediators (1, 3-5). Notably, defects in plasmalogen biosynthesis have been linked to severe neurological pathologies, including Zellweger syndrome (6) and rhizomelic chondrodysplasia punctata (1, 7). Early investigations proposed that plasmalogens act as sacrificial antioxidants based on the high reactivity of their vinyl-ether groups with reactive oxygen species, thereby shielding adjacent PUFA from oxidation (8-11). However, recent findings have challenged this notion by revealing a prooxidant and pro-ferroptotic role for ether phospholipids (12-15). For example, it has been found that elevated plasmalogen levels in cardiomyocytes and neurons enhance their susceptibility to ferroptosis (13). However, the mechanisms through which plasmalogens influence cell sensitivity to lipid oxidation remains elusive.

Singlet molecular oxygen [O_2_ (^1^Δ_g_)] is of great importance in chemical and biological systems due to its high reactivity and involvement in physiological and pathological processes (16). Additionally, it serves as tumoricidal and antibacterial component in photodynamic therapy (17). O_2_ (^1^Δ_g_) is generated by light-dependent photosensitized reactions and by light-independent chemical and biochemical reactions (18, 19). Singlet oxygen exhibits substantial reactivity towards electron-rich biological molecules, including DNA, proteins, and lipids (18). Among membrane components, plasmalogen vinyl ether bond is highly vulnerable to singlet oxygen mediated oxidation. The rate constants for both physical and chemical quenching of O_2_ (^1^Δ_g_) by vinyl-ether groups have been estimated to be in the range of 10^6^-10^7^ M^−1^s^−1^ (20). This is notably 1-2 orders of magnitude faster compared to the reaction with diacyl phospholipid analogues, indicating that plasmalogens are among the primary targets for O_2_ (^1^Δ_g_) oxidation in biological membranes. However, a comprehensive characterization of primary and secondary oxidation products of plasmalogens is still lacking.

Previous studies have described the generation of dioxetane and hydroperoxide intermediates in plasmalogen oxidation (10, 21, 22). These intermediates can act as prooxidant species by generating excited molecules and free radicals. For example, the thermolysis of dioxetane produces highly reactive triplet excited carbonyl species (18, 23), which subsequently decay to the ground state emitting light in the visible region or transfer energy to molecular oxygen, resulting in the production O_2_ (^1^Δ_g_) (23, 24) (Fig 1). Notably, the production of chemiexcited species has attracted attention in recent years due to its implications in skin cancer and eye diseases (25-27). Additionally, lipid hydroperoxides can react with metal ions, heme proteins, or other oxidants, producing alkoxyl and peroxyl radicals (28). These radicals can further propagate lipid peroxidation by fueling radical chain reactions and by promoting the generation of excited species, such as O_2_ (^1^Δ_g_) (29-33).

**Fig. 1.**
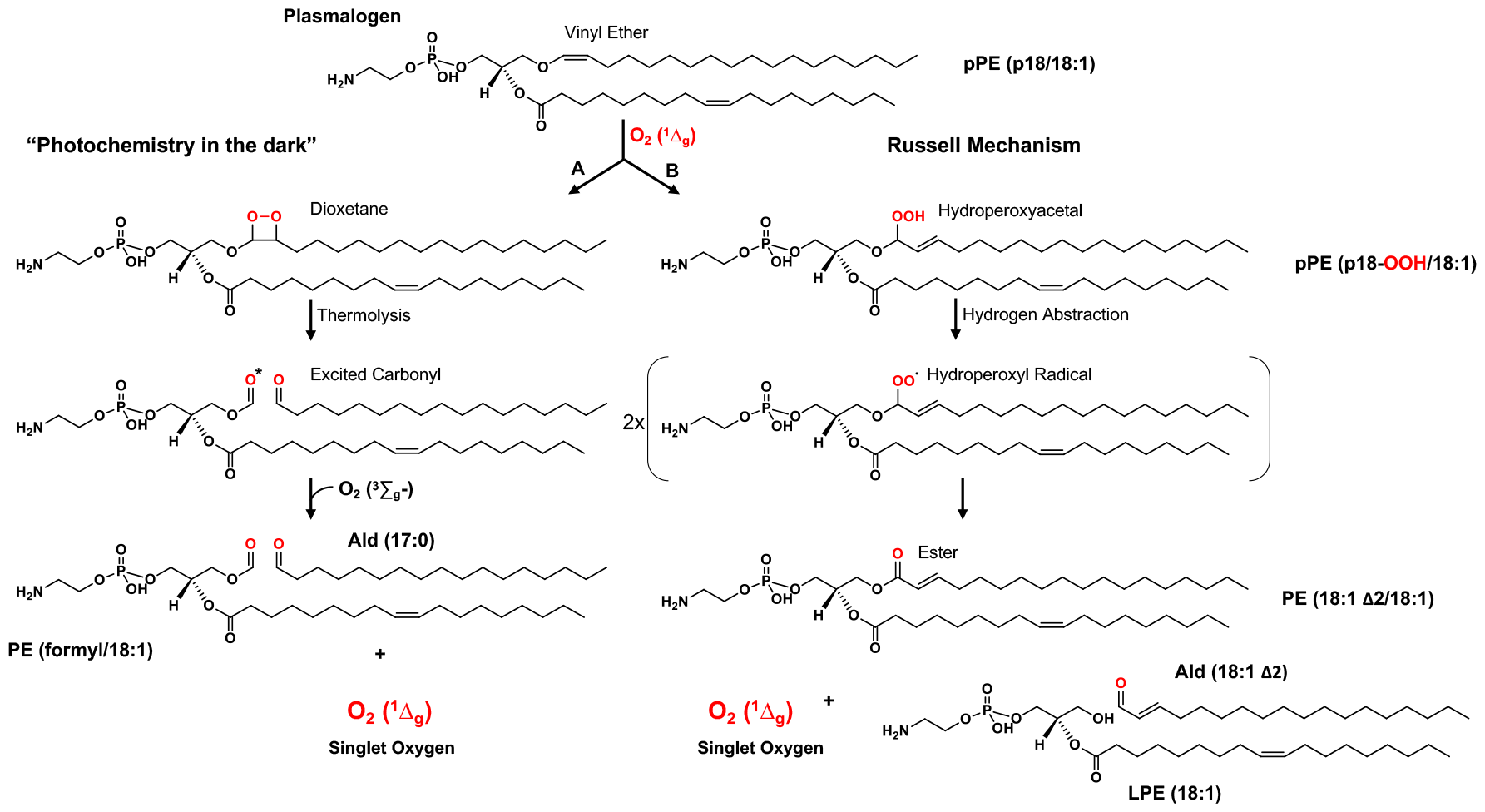
Scheme depicting the oxidation of plasmalogen vinyl ether bond by singlet oxygen. pPE (p18:0/18:1) undergoes reaction with O2 (1Δg), resulting in the formation of dioxetane and hydroperoxylacetal intermediates. (A) Thermal decomposition of 1,2-dioxetane generates carbonyl compounds, including one in a triplet-excited state (*). These high-energy triplet-excited carbonyls can transfer energy to molecular oxygen, leading to the production of O_2_ (^1^Δ_g_). Subsequent C-C bond rupture yields a formyl-PE and a fatty aldehyde. (B) Decomposition of hydroperoxyacetal results in the formation of peroxyl radicals, either through H-atom transfer or metal-catalysis. These peroxyl radicals then react, producing O_2_ (^1^Δ_g_), diacyl PE, lysoPE (LPE), and an alpha-beta-unsaturated fatty aldehyde.

In this study, our aim was to thoroughly investigate the formation of reactive species resulting from plasmalogen oxidation with O_2_ (^1^Δ_g_). By utilizing chemiluminescence and high-resolution mass spectrometry analysis to characterize primary and secondary oxidation products, we demonstrate the generation of prooxidant and electrophilic lipid species. Our finding reveals that the plasmalogen vinyl-ether bond reacts with O_2_ (^1^Δ_g_), leading to the generation of hydroperoxy acetal at the sn-1 position as the major primary oxidation product. Furthermore, we provide evidence for the generation of excited species through chemiluminescence and chemical trapping analysis. Additionally, we developed a specific LC-MS method to characterize the production of electrophilic lipid species, including alpha-beta unsaturated phospholipids and long-chain fatty aldehydes. Together, our study highlights the generation of oxidizing excited molecules and reactive lipid species from plasmalogens.

## Results

### Characterization of plasmalogen photooxidation products

To characterize plasmalogen photooxidation products, a sample of pPE (p18:0/18:1) was photooxidized at low temperature (-40°C). This procedure aimed to prevent dioxetane decomposition. Aliquots of the sample were collected at various time points, and the oxidation products were analyzed by liquid chromatography coupled to high-resolution mass spectrometry (LC-Q-TOF-MS/MS). During photooxidation, the pPE (p18/18:1) peak at 10.3 minutes was rapidly consumed, nearly depleting after 30 minutes. Notably, plasmalogen consumption was accompanied by the appearance of a major peak with higher polarity at 9.6 min, along with some minor polar products eluting at earlier retention times, e.g., 4.3 min (**Fig. 2A**). The photooxidation products were identified through MS (**Fig. 2B**) and MS/MS analysis (**Fig. 2C-F**). As expected, pPE (p18:0/18:1) exhibited the deprotonated ion [M-H]^−^ at m/z 728.5601 (**Fig. 2B**) and typical fragment ions corresponding to the phosphoethanolamine head-group (m/z 140.0118), the 18:1 acyl chain (m/z 281.2486), and two additional fragments resulting from 18:1 acyl chain loss as an acid (m/z 446.3023) or a ketene (m/z 464.3146) (**Fig. 2C**).

**Fig. 2.**
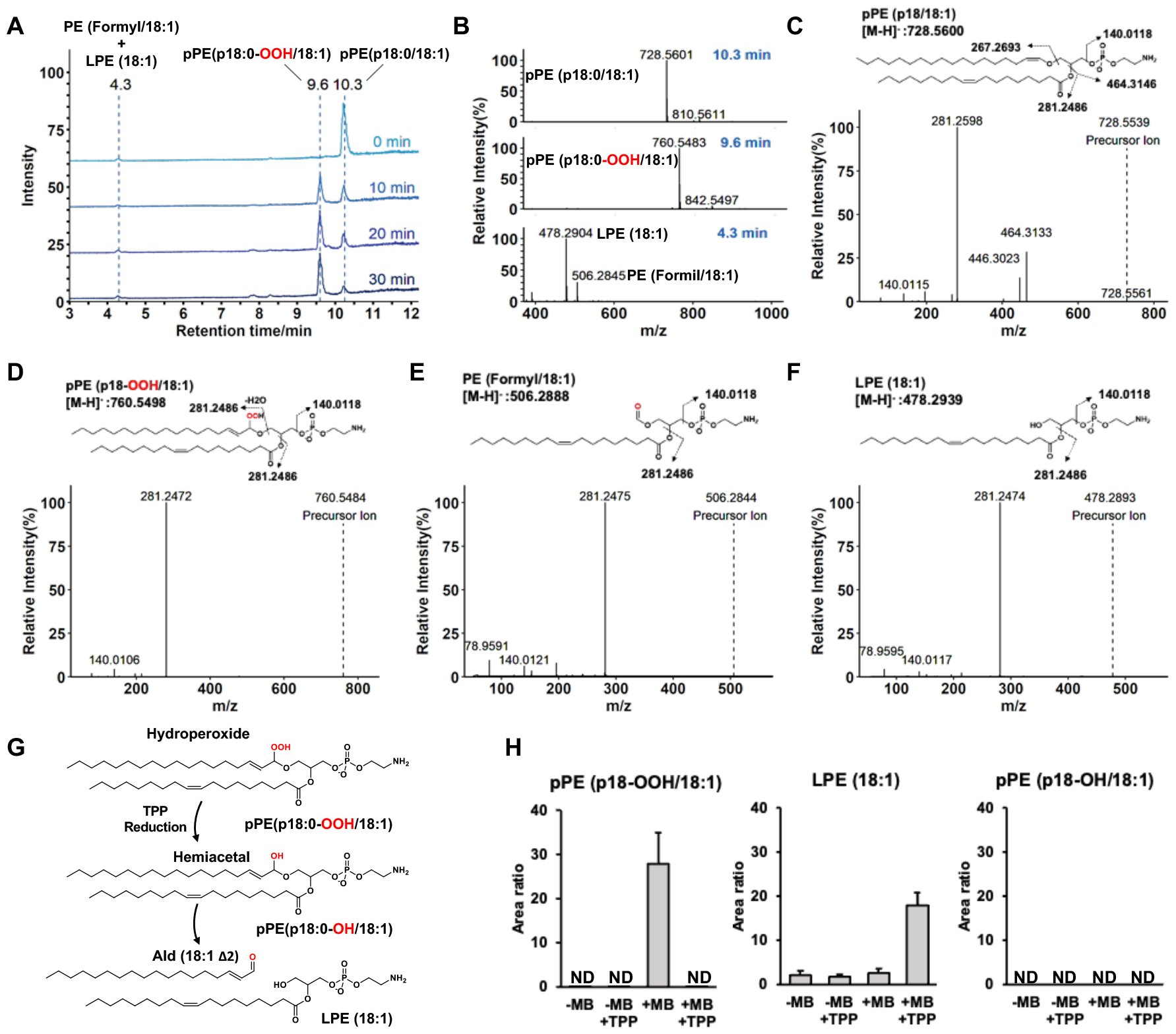
Characterization of plasmalogen photooxidation products by LC-QTOF-MS analysis. (A) Total ion chromatograms of pPE (p18:0/18:1) photooxized with MB for 0, 10, 20 and 30 min. (B) MS1 mass spectra of pPE oxidation products detected at 4.3, 9.6 and 10.3 min. (C-F) MS2 mass spectra of pPE (p18:0/18:1); pPE(p18:0-OOH:18:1); LPE(18:1); and PE (formyl/18:1). (G) Reaction scheme illustrating the reduction of pPE (p18-OOH/18:1) with triphenylphosphine (TPP), resulting in the formation of a hemiacetal. This hemiacetal undergoes cleavage, yielding a fatty aldehyde, Ald (18:1Δ2), and LPE (18:1). (H) Relative concentrations of plasmalogen oxidized products before and after TPP addition. Reaction conditions: 1 mM pPE (p18/18:1) was photooxidized in the presence of 50 μM methylene blue at -40 °C under the following conditions: (-MB) without methylene blue; (+MB) with methylene blue; with MB and with TPP (+MB + TPP). Data are shown as area ratio ± SEM

The major product detected at 9.6 minutes exhibited an m/z value at 760.5483 consistent with two oxygen atom addition to the vinyl ether bond. This product was identified as pPE(p18:0-OOH/18:1), a plasmalogen containing hydroperoxyacetal group at sn1 position. The MS/MS spectra of this product revealed a prominent fragment ion at m/z 281.2486, originating from both the dissociation of the sn2-18:1 acyl chain and the dissociation and dehydration (-18) of the sn1 hydroperoxide alkyl chain (**Fig. 2D**). Furthermore, to confirm the identity of hydroperoxiacetal group we used triphenylphosphine (TPP) as a reducing agent. Hydroperoxiacetal reduction is expected to produce a hemiacetal intermediate that is cleaved yielding lysophosphatidylethanolamine [LPE (18:1)] and an alpha-beta unsaturated fatty aldehyde [Ald (18:1 Δ2)] (**Fig. 2G**). Indeed, TPP reduction led to a huge increase in LPE (18:1), confirming the hydroperoxyacetal group (**Fig. 2H** and **Fig. S3**). Of note, pPE hydroxide was not detected, thus excluding the possibility of a hydroxide group being formed in the 18:1 acyl chain. Besides the plasmalogen hydroperoxyacetal peak at 9.6 min, two minor polar products eluting at around 4.3 min were identified as LPE (18:1) and PE (formyl/18:1). These products had similar MS/MS spectra showing major fragments at m/z 281.247 and 140.012 corresponding to the 18:1 acyl chain and phosphoethanolamine, respectively. However, PE (formyl/18:1) MS spectra showed a mass shift of +27.99 Da due to the formyl group (**Fig. 2E and F**).

To further investigate the formation of plasmalogen sn-1 hydroperoxyacetal in a more complex sample, a plasmalogen sample purified from bovine brain was photooxidized under the same reaction conditions (**Fig. S1**). Interestingly, all plasmalogen species were rapidly consumed within 30 min of reaction, yielding the corresponding hydroperoxacetal products (**Table S1**). The formation hydroperoxide at the vinyl-ether bond in plasmalogen was confirmed by the MS/MS analysis. All MS spectra showed the characteristic sn2 fatty acyl chain fragment ions and the sn-1 alkyl chain containing the hydroperoxyacetal group, which showed a mass shift of -18 due to the dehydration of the hydroperoxide group (**Fig. S2**). Furthermore, reduction with TPP resulted hydroperoxide consumption leading to a huge increase in the corresponding LPE products (**Table S1**). Thus, these data demonstrates that plasmalogen oxidation by O_2_ (^1^Δ_g_) generates hydroperoxyacetal as major primary products.

### Determination of the ratio between plasmalogen photooxidation routes

The reaction of singlet oxygen with the plasmalogen vinyl ether group can proceed through two pathways (Fig.1). The first pathway involves a [2+2] cycloaddition reaction mechanism, leading to the formation of 1,2-dioxetanes and the second pathway involves an *ene* reaction mechanism, resulting in hydroperoxides. Dioxetanes thermally degrade, giving rise to formyl-PE and fatty aldehydes (Route A), while ether-hydroperoxides can be converted to a hemiacetal intermediate, which spontaneously cleaves, producing LPE and fatty aldehydes (Route B).

To get deeper insights into the mechanism of plasmalogen oxidation via singlet oxygen reaction, we determined the relative amounts of oxidized phospholipid products and long-chain aldehydes arising from both pathways by LC-MS analysis. Among the oxidized phospholipid products, PE (formyl/18:1) and pPE (p18:0-OOH/18:1), were used as markers for dioxetane and hydroperoxide pathways, respectively. Notably, plasmalogen hydroperoxyacetal was detected as major product at all time points of plasmalogen photooxidation (**Fig. 3B**). The hydroperoxide-to-dioxetane ratio estimated by this approach was 98:2. This ratio was further corroborated by the quantification of long-chain fatty aldehydes, Ald (17:0) and Ald (18:1Δ2), generated from dioxetane and hydroperoxide, respectively (**Fig. 3A**). Both fatty aldehydes were detected at very low levels in photooxidized samples. In contrast, Ald (18:1Δ2) levels strikingly increased after TPP reduction (**Fig.3C**). The relative ratio fatty aldehydes yielded an estimated hydroperoxide-to-dioxetane ratio of 96:4, closely matching the value obtained in the phospholipid analysis. Together this data show that vinyl ether bond oxidation by O_2_ (^1^Δ_g_) predominantly proceeds the *ene* reaction mechanism, yielding plasmalogen hydroperoxyacetal as a major primary product.

**Fig. 3.**
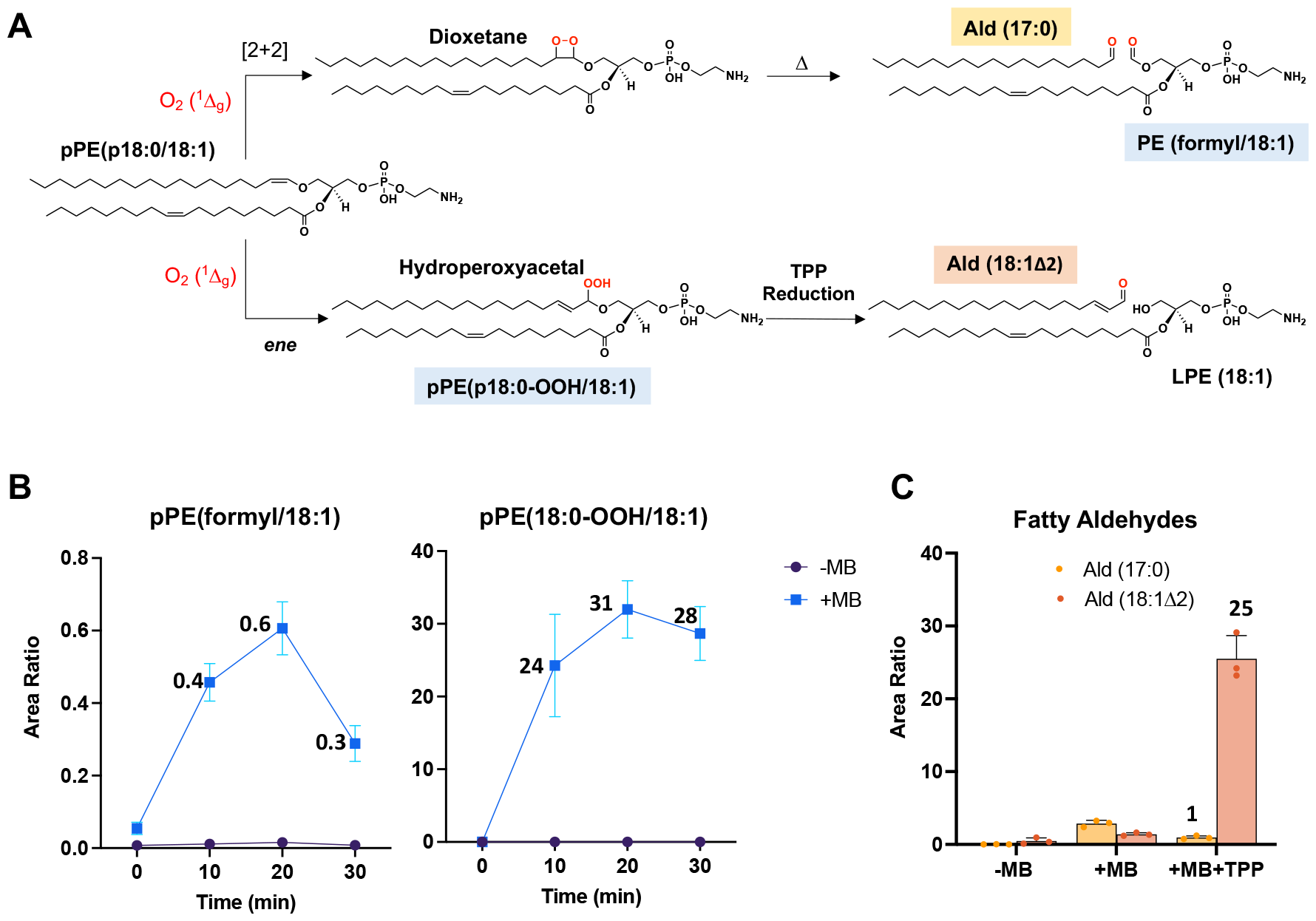
Determination of the relative ratio between dioxetane and hydroperoxyacetal pathways. (A) Reaction scheme illustrating the oxidation of plasmalogen vinyl ether bond by O_2_ (^1^Δ_g_). Top-The [2+2] cycloaddition mechanism leads to the formation of dioxetane, which undergoes subsequent cleavage to produce PE (formyl/18:1) and Ald (17:0). Bellow-The ene-reaction generates hydroperoxyacetal, which is then reduced to produce LPE (18:1) and Ald (18:1Δ2). (B) Quantification of PE (formyl/18:1) and pPE (p18:0-OOH/18:1). (B) Quantification of fatty aldehydes, Ald (17:0) and Ald (18:1Δ2) after 30 minutes of irradiation. Reaction conditions were the same as in Figure 2. Data are shown as area ratio±SEM.

### Characterization of fatty aldehydes generated from plasmalogen hydroperoxides

Previous studies have detected long-chain fatty aldehydes from plasmalogen oxidation (21, 34-37). Here we developed a LC-MS method to specifically detect fatty aldehydes using the 7-(diethylamino)coumarin-3-carbohydrazide (CHH) (38) and 7-mercapto-4-methylcoumarin (CSH) probes. As mentioned above, heptadecanal [Ald (17:0)] was used as a marker for dioxetane and 2-octadecenal [Ald (18:1 Δ2)], an alpha-beta unsaturated fatty aldehyde, was used as a marker for plasmalogen hydroperoxides (pPE (p18-OOH/18:1)).

Firstly, aldehydes derivatized with the CHH probe **(Fig. 4A)** were characterized by mass spectrometry. The Ald (17:0)-CHH adduct detected in photooxidized pPE exhibited the same retention time and fragmentation profile as the commercial Ald (17:0) standard. Both MS/MS spectra showed a prominent fragment ion corresponding to the CHH probe at m/z 244.0968 and a fragment ion corresponding to the fatty alkyl chain at m/z 252.2702 (**Fig. 4B and C**). Similarly, Ald (18:1 Δ2)-CHH adduct, showed a precursor ion at m/z 524.3847, and a fragment ion corresponding to the fatty alkyl chain at m/z 279.2830 (**Fig. 4D**).

**Fig. 4.**
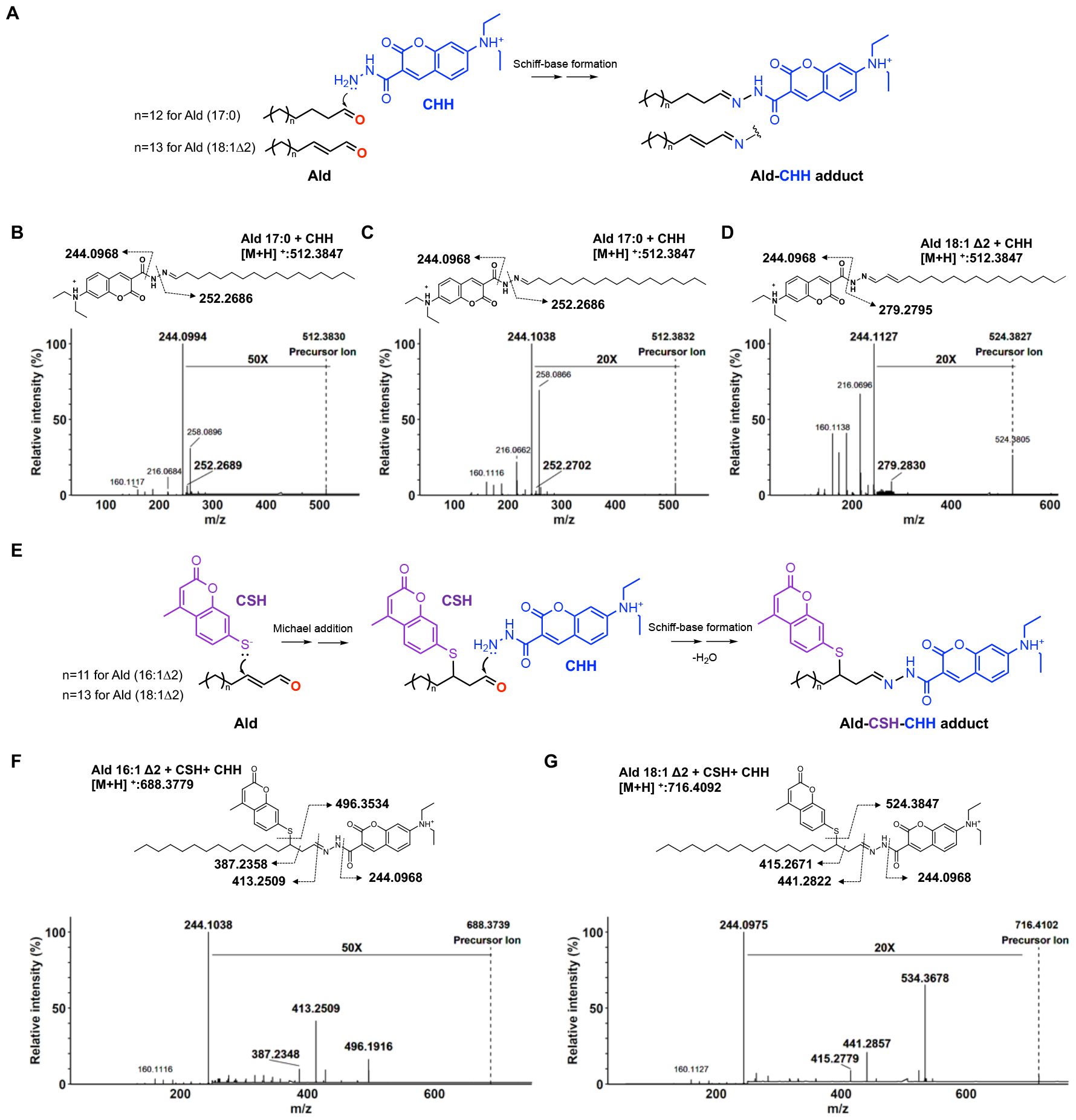
Characterization of long-chain fatty aldehydes by mass spectrometry. (A) Scheme illustrating fatty aldehyde (Ald) derivatization with the CHH probe. MS/MS spectra of the Ald-CHH adduct from (B) a commercial standard of Ald (17:0); (C) Ald (17:0) generated from pPE (p18/18:1) photooxidation; and (D) Ald (18:1 Δ2) generated from pPE (p18:0-OOH/18:1) reduction with TPP. (E) Scheme depicting alpha-beta unsaturated fatty aldehyde derivatization with CSH and CHH probes. MS/MS spectra of the Ald-CSH-CHH adduct from (F) a commercial standard of Ald (16:1 Δ2); and (G) Ald (18:1 Δ2) generated from pPE (p18:0-OOH/18:1) reduction with TPP.

Next, fatty aldehydes containing alpha-beta carbonyl group was derivatized with CSH, a coumarin derivative containing a thiol group, as a second probe. The derivatization method using the CHH and CSH fluorescent probes was validated using commercially available Ald (16:1 Δ2) and Ald (16:0) (**Fig. S4)**. As illustrated for Ald (16:1 Δ2) the thiol group in the CSH probe reacts with the beta-carbon in the fatty aldehyde, preserving the aldehyde group and allowing subsequent derivatization with the CHH probe **(Fig. 4E**). The doubly derivatized aldehyde exhibited a fragmentation profile showing the loss of the probes and fragments indicating the presence of the alpha-beta unsaturated carbonyl group (**Fig. 4F**). Of note, a similar fragmentation pattern was observed for Ald (18:1 Δ2) detected in the photooxidized plasmalogen sample after TPP reduction (**Fig. 4G**), clearly confirming the presence of alpha-beta unsaturated carbonyl groups.

### Detection of excited triplet carbonyls and singlet oxygen from oxidized plasmalogen species

Dioxetane decomposition generates excited triplet carbonyls and O_2_ (^1^Δ_g_) (39, 40). Excited triplet carbonyls have been linked to normal and deleterious effects due to their radical-like chemical and physicochemical properties (41, 42). They often return to the ground state producing a ultraweak photon emission in the visible region (43) and participate in energy transfer reactions with other molecules, including molecular oxygen [O_2_ (^3^Σ_g_)], producing O_2_ (^1^Δ_g_) (40)(**Fig. 5A**).

**Fig. 5.**
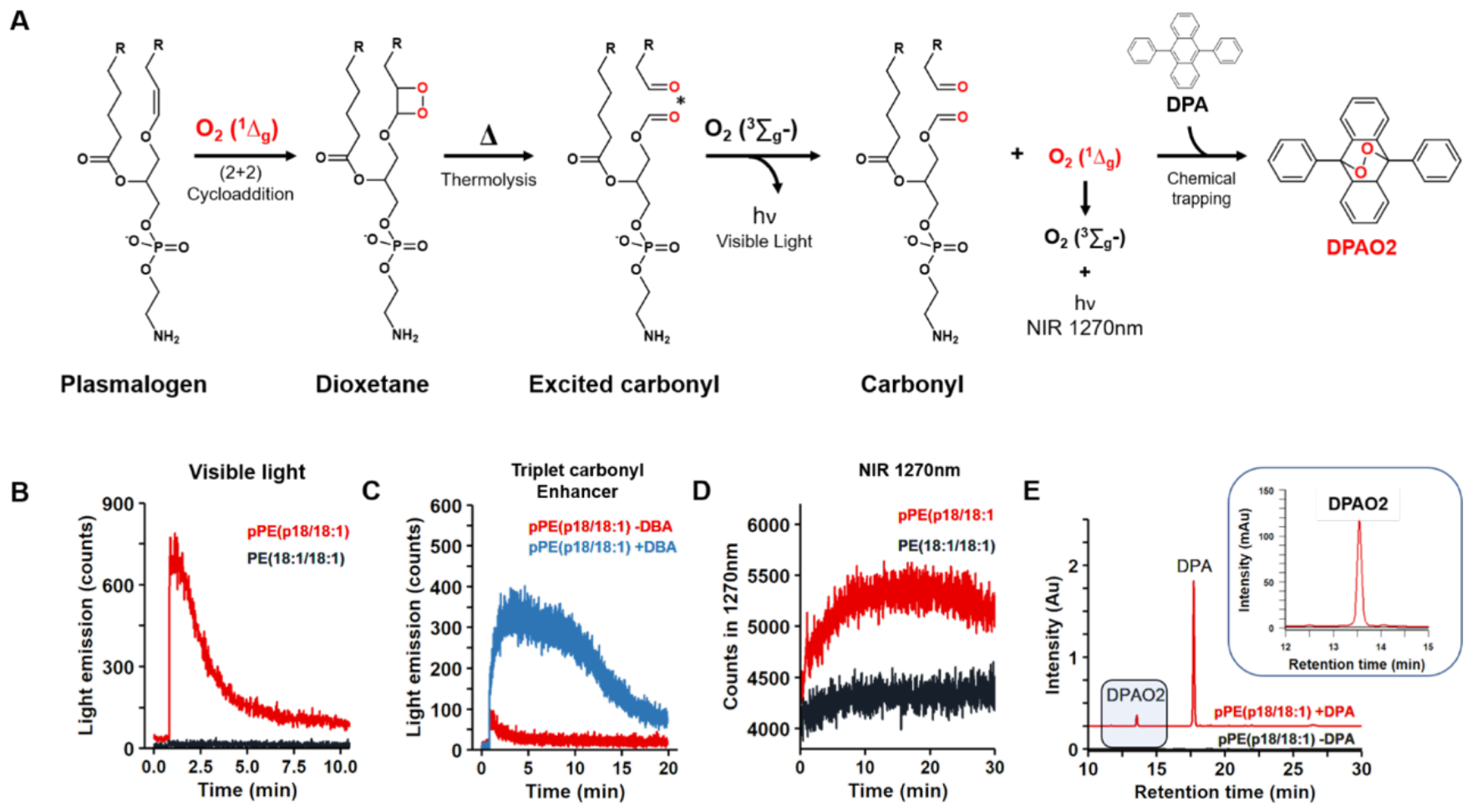
Generation of excited triplet carbonyls and singlet oxygen from plasmalogen dioxetane intermediate. (A) Scheme illustrates the formation of dioxetane through the reaction of O_2_ (^1^Δ_g_) with the vinyl-ether bond. Dioxetane thermolysis yields carbonyl species, one of them in the excited state. These excited triplet carbonyls can emit light in the visible region or transfer energy to molecular oxygen, resulting in the generation of O_2_ (^1^Δ_g_). Singlet oxygen production was confirmed by light emission in the NIR region (1270 nm) and by chemical trapping of O_2_ (^1^Δ_g_) with DPA. (B) Visible light emission resulting from dioxetane thermolysis (red). (C) Enhancement of triplet carbonyl light emission using 10 mM DBA (blue). (D) Near-infrared (NIR) light emission at 1270 nm (red). (E) Chemical trapping of O_2_ (^1^Δ_g_) by DPA and analysis of DPAO2 by HPLC (UV 210 nm). Light emission and chemical trapping experiments were performed using 1 mM pPE (p18/18:1) (red lines) or PE (18:1/18:1) (black lines) solutions that had been photooxidized with MB at -40°C for 30 minutes.

To demonstrate the generation of excited triplet carbonyls, plasmalogen samples photooxidized at -40°C were allowed to naturally heat up to room temperature, and visible light emission was recorded using a photomultiplier. As expected, intense light emission was observed from the photooxidized pPE (p18/18:1) sample but not from PE (18:1/18:1) (**Fig. 5B**).The generation of triplet excited carbonyls was further confirmed by using dibromoanthracene (DBA) as a triplet carbonyl enhancer (41), which produced a noticeable enhancement in the visible light emission (**Fig. 5C**).

After detecting triplet carbonyls, our subsequent investigation focused on the detection of O_2_ (^1^Δ_g_) generated through energy transfer from excited triplet carbonyls to molecular oxygen. O_2_ (^1^Δ_g_) detection was performed by the measurement of its monomolecular light emission in the near-infrared (NIR) region at 1270 nm (18, 30). Notably, a light emission persisting for several minutes was observed in the photooxidized pPE (p18:0/18:1) (**Fig. 5D**). The production of O_2_ (^1^Δ_g_) was confirmed by chemical trapping with 9,10-diphenylanthracene (DPA). This chemical probe reacts rapidly (k = 1.3 X10^6^ M^-1^s^-1^) with O_2_ (^1^Δ_g_) resulting in a stable endoperoxide (DPAO_2_) that can be detected by HPLC (30, 44) (**Fig. 5A**). DPA was added after photooxidation at low temperature and incubated at room temperature for 1 hour in the dark. HPLC analysis confirmed the formation of DPAO_2_ (**Fig. 5E**). Together, these data confirm the production of O_2_ (^1^Δ_g_) from photooxidized plasmalogen species.

### Characterization of plasmalogen hydroperoxyacetal decomposition products

Lipid hydroperoxides react with metal ions such as FeII and CeIV, producing peroxyl radicals and O_2_ (^1^Δ_g_) via the Russell mechanism (30, 45). In the Russell mechanism two peroxyl radicals reacts head-to-head to form a linear tetroxide intermediate that decomposes producing a ketone, alcohol and O_2_ (^1^Δ_g_) (29). We have previously demonstrated that lipid hydroperoxides generates O_2_ (^1^Δ_g_) and the corresponding alcohol and ketone products through the Russell mechanism (30-32, 46). Here, our aim was to characterize the major oxidation products resulting from the metal ion-catalyzed decomposition of pPE (p18:0-OOH/18:1). Plasmalogen hydroperoxyacetal reaction with metal ions produces peroxyl radicals that produces a hemiacetal and a ketone as final products. The hemiacetal intermediate is cleaved, producing LPE (18:1) and Ald (18:1 Δ2), while the ketone group forms an ester bond at sn1 position producing a diacyl-PE, (**Fig. 6A**). As a comparison we also studied the products formed by the reaction of PE hydroperoxides (**Fig. 6B**).

**Fig. 6.**
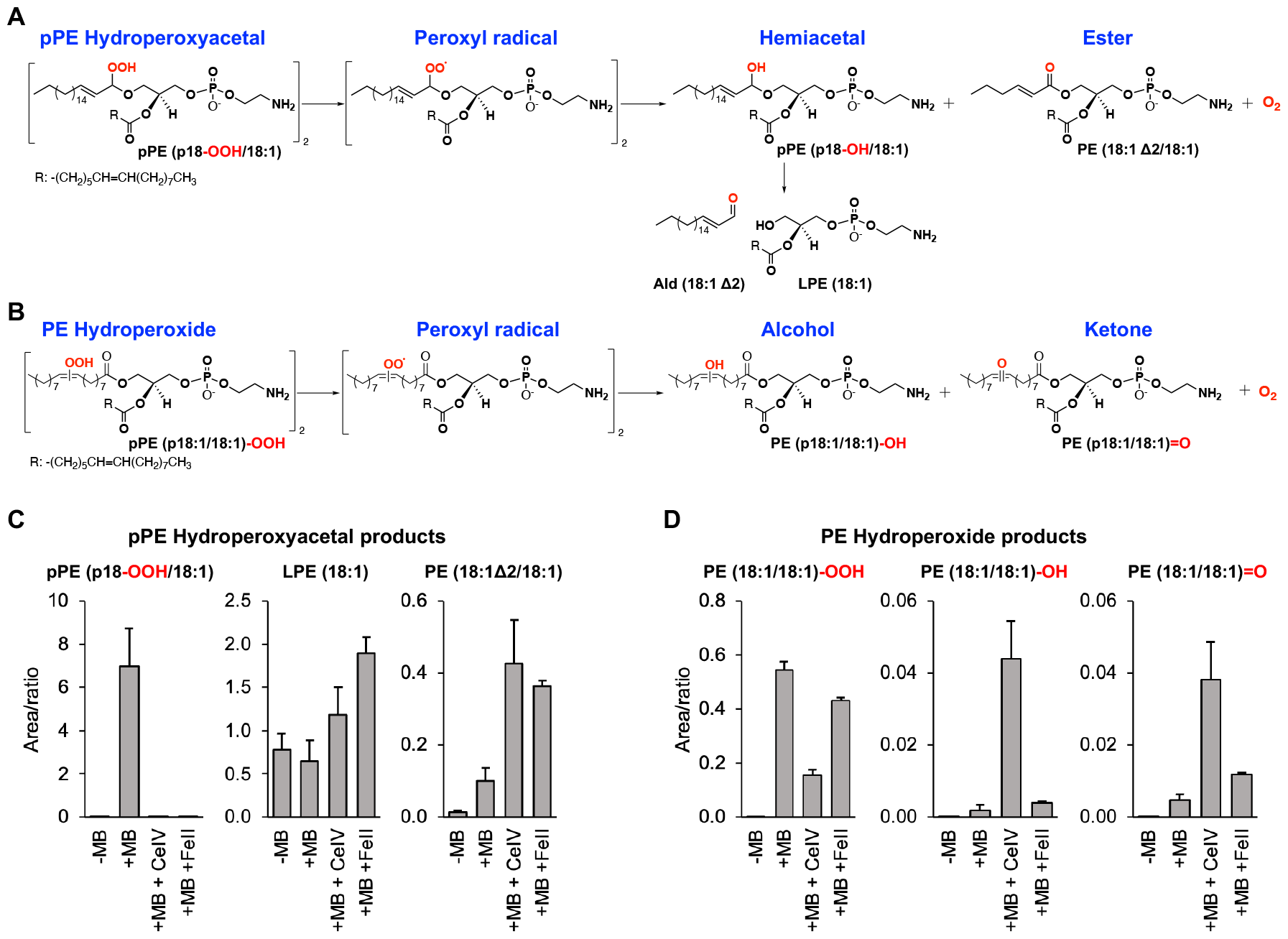
Plasmalogen hydroperoxyacetal reacts with metal ions, producing alpha-beta unsaturated fatty aldehydes and lyso-PE. (A-B) Scheme depicting the reactions products for pPE-hydroperoxyacetal and PE-hydroperoxides. (C-D) Quantification of phospholipid oxidation products by LC-QTOF-MS. Reaction conditions: A solution of 1 mM plasmalogen solution photooxidized with MB at -40C for 30 min were mixed with 50 μM CeIV of FeII in CHCl_3_:methanol:water (90:9:1 v/v/v) for 1 hour.

Mass spectrometry analysis showed that plasmalogen hydroperoxyacetal are completely consumed in the presence of CeIV or FeII, producing LPE (18:1) and PE (18:1Δ2/18:1) as major products (**Fig. 6C**). In contrast, some PE hydroperoxide remained intact after incubation with CeIV or FeII. As expected, PE hydroperoxide decomposition generated PE species containing the corresponding alcohol and ketone groups (**Fig. 6D**). Together these results shows that plasmalogen hydroperoxyacetal is totally decomposed by metal ions producing LPE and an ester, the diacyl-PE containing alpha-beta unsaturated carbonyl group at the sn-1 position. The diacyl PE showed the same MS/MS spectrum of PE (18:1/18:1) (**Fig. S5**), albeit with slightly different retention times probably due to different location of the double bond in the sn1 acyl chain (**Fig. 7A**). Furthermore, we confirmed the presence of alpha-beta unsaturated carbonyl group by reacting with the CSH probe and detecting PE (18:1Δ2/18:1)-CSH adduct by mass spectrometry. The MS/MS spectra of this adduct showed the characteristic CSH fragment ion at m/z 191.0172, along with fragment ions arising from sn1 and sn2 acyl chain cleavages (**Fig. 7B**).

**Fig. 7.**
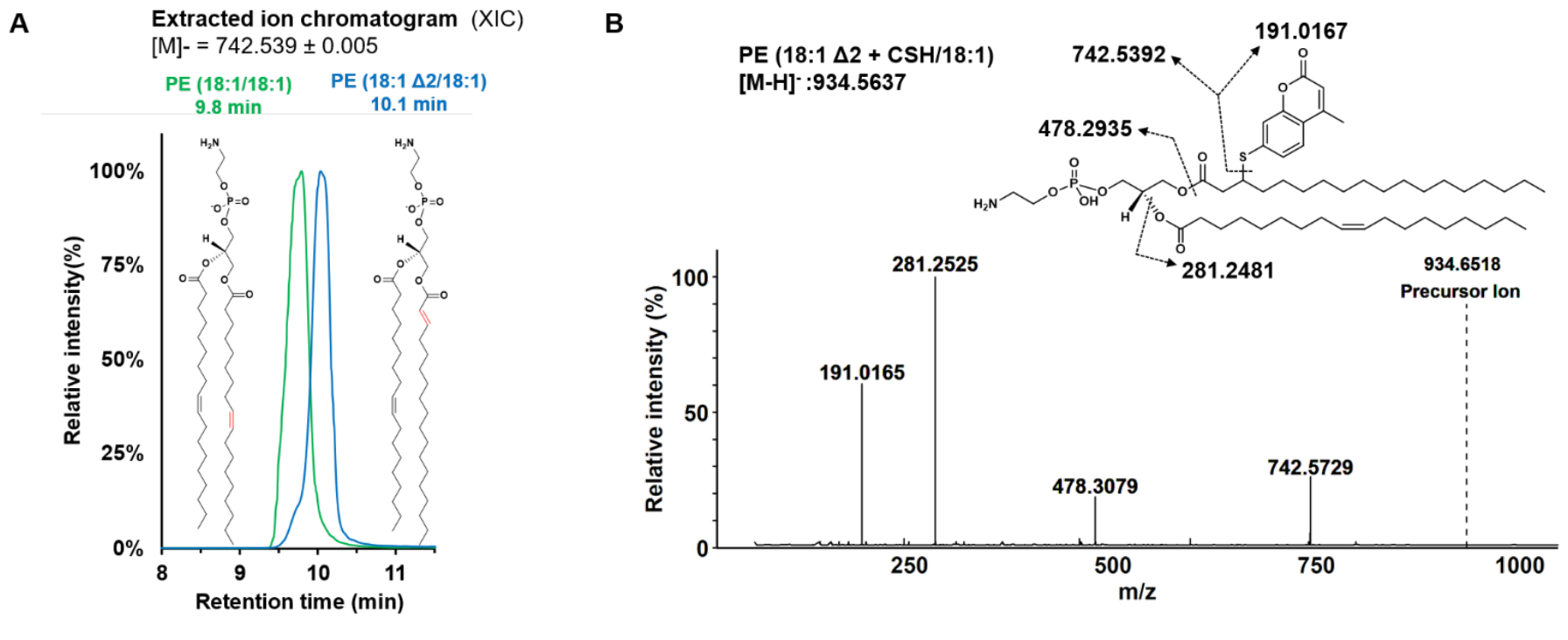
Characterization of alpha-beta unsaturated carbonyl group in PE (18:1Δ2/18:1). (A) Extracted ion chromatogram of m/z 742.539 in photooxidized pPE(p18/18:1) treated with CeIV (blue) and the commercial PE(18:1/18:1) standard (green). (B) MS/MS spectrum of PE(18:1Δ2/18:1)-CSH adduct.

### Characterization of singlet molecular oxygen production from plasmalogen hydroperoxyacetal

To examine the production of O_2_(^1^Δ_g_) resulting from the reaction of plasmalogen hydroperoxyacetal with metal ions, we measured the light emission in the NIR region at 1270 nm. An intense emission signal was observed in the reaction of pPE (p18:0-OOH/18:1) with CeIV (**Fig. 8A**). For comparison, we also measured the light emission generated from linoleic acid hydroperoxide (LAOOH) (**Fig. 8B**), a positive control that has been previously shown to produced O_2_(^1^Δ_g_) via the Russell’s mechanism (30). Interestingly, both hydroperoxides produced light emission at 1270 nm with similar intensity and kinetics.

**Fig. 8.**
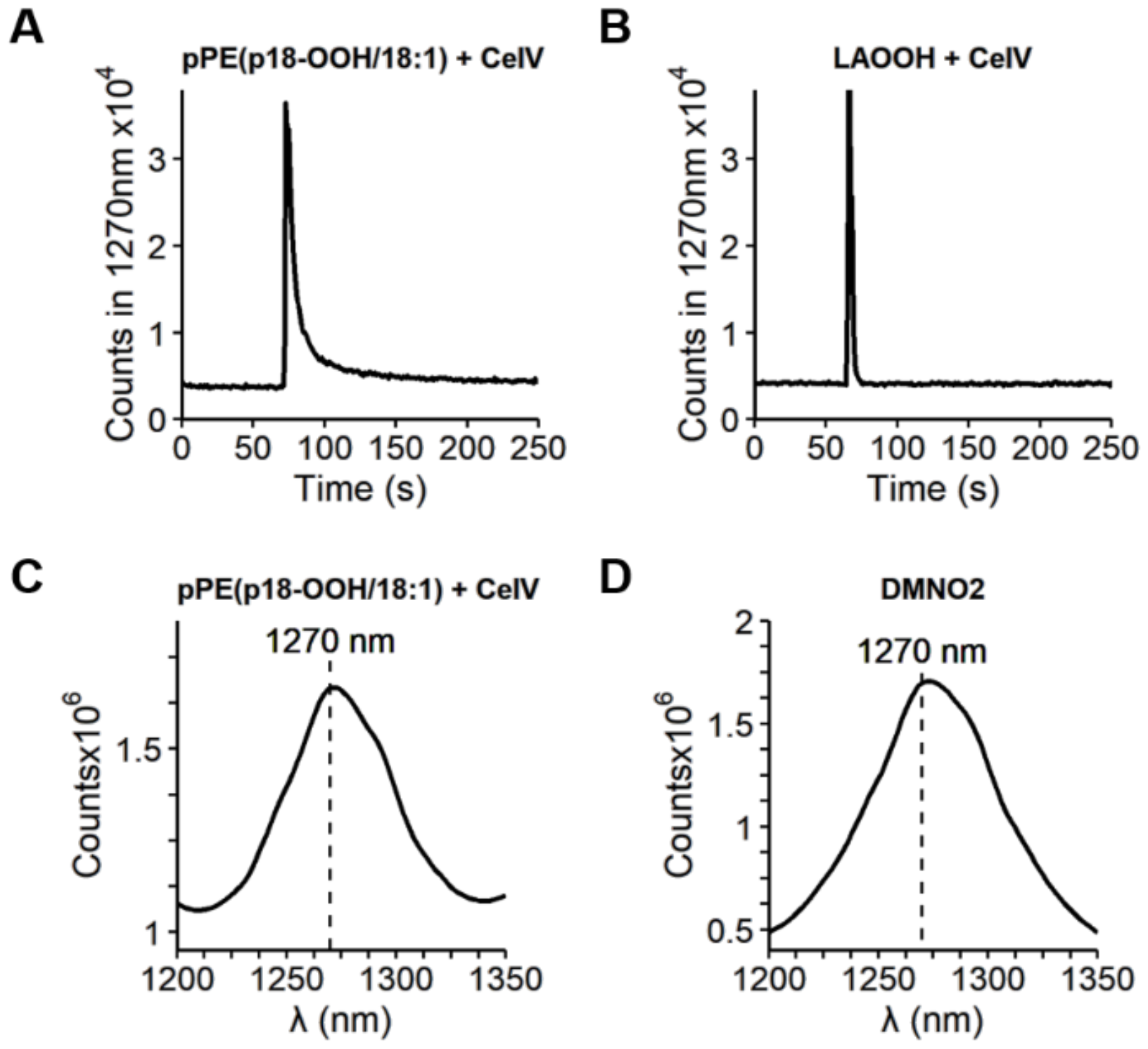
Singlet oxygen generation from plasmalogen hydroperoxyacetal reaction with CeIV. (A) Light emission signal at 1270 nm obtained for the reaction of 5 μM pPE (p18:0-OOH/18:1). (B) Comparison with the reaction using linoleic acid hydroperoxide (LAOOH). (C) NIR light emission spectrum obtained in the reaction of 100 μM pPE (p18:0-OOH/18:1) with 75 μM CeIV. (D) NIR light emission spectrum obtained during the thermolysis of 1 mM DMNO2.

Furthermore, we confirmed the generation of O_2_ (^1^Δ_g_) from pPE (p18:0-OOH/18:1) by the acquisition of the NIR light emission spectrum, showing a maximum intensity at 1270 nm (**Fig. 8C**). As a control, we also acquired the emission spectra for DMNO_2_, a classical chemical generator of O_2_ (^1^Δ_g_) (**Fig. 8D**). To estimate the relative yield of O_2_(^1^Δ_g_) we used the emission signal from DMNO2 as a reference (**Fig. S6**). The yield of O_2_(^1^Δ_g_) generated from plasmalogen hydroperoxide was 3.2%, consistent with previous yields of O_2_(^1^Δ_g_) reported for lipid hydroperoxides reacting by the Russell’s mechanism (30). Together, these results clearly demonstrate that plasmalogen hydroperoxyacetal generate O_2_(^1^Δ_g_) upon reaction with metal ions.

## Discussion

In this study, we demonstrate that the oxidation of plasmalogens by O_2_ (^1^Δ_g_) generates reactive intermediates capable of promoting the formation of excited species and electrophilic lipid species. Chemiluminescence measurements conducted in the visible and near-infrared regions (specifically at 1270 nm) detected characteristic light emissions associated with the production of excited triplet carbonyls and O_2_ (^1^Δ_g_). The generation of excited triplet carbonyls from dioxetane was confirmed through light emission measurements in the visible region, as well as by observing enhanced light emission upon the addition of DBA, a triplet chemiluminescence enhancer. Moreover, our study offers compelling evidence demonstrating the production of O_2_ (^1^Δ_g_) through the reaction between plasmalogen hydroperoxyacetal and metal ions. Particularly, the intense light emission signal detected in the near-infrared region, showing an emission peak at 1270 nm, serve as unequivocal evidence for O_2_ (^1^Δ_g_) generation.

In terms of mechanism, the generation of excited triplet carbonyls and O_2_ (^1^Δ_g_) from plasmalogen oxidation can be explained by two pathways as depicted in Figure 1. Firstly, the [2 + 2] cycloaddition results in the formation of an unstable 1,2-dioxetane intermediate, which then undergoes thermolysis to produce excited triplet carbonyls (Fig. 1A). Secondly, the *ene* reaction leads to the formation of plasmalogen hydroperoxyacetal. This compound subsequently generates O_2_ (^1^Δ_g_) through reactions involving the combination of peroxyl radicals via the Russell mechanism (Fig. 1B).

The relative proportion between these two pathways was carefully examined using mass spectrometry analysis of phospholipid oxidation products. Our data reveals that plasmalogen predominantly reacts with O_2_ (^1^Δ_g_) through the *ene* reaction, yielding plasmalogen hydroperoxyacetal as the major product (97%), while the formyl-PE resulting from dioxetane cleavage was detected at minimal quantities (3%). This finding was further supported by the analysis of relative yields of fatty aldehydes derived from dioxetane and hydroperoxide intermediates. We employed an LC-MS method specifically developed to detect fatty aldehydes containing alpha-beta unsaturated carbonyl groups. Notably, Ald (18:1 Δ2), the alpha-beta unsaturated fatty aldehyde derived from plasmalogen PE (p18:0-OOH/18:1), significantly increased after reduction with TPP, becoming the most abundant aldehyde (96%) detected in the reaction. In contrast, heptadecanal (Ald 17:0), a product formed by dioxetane cleavage, was detected at minor levels (4%). Thus, we confirmed using two different LC-MS methods that plasmalogen photooxidation predominantly generates hydroperoxyacetal as the major primary photooxidation product.

Earlier research has reported the detection of plasmalogen hydroperoxyacetal, which forms at the vinyl ether bond of plasmalogen species through photosensitized and radical oxidations (10, 21, 22). Although the biological fate of these hydroperoxides remains unknown, it is well established that phospholipid hydroperoxides are rapidly reduced to hydroxides by antioxidant enzymes, such as GPx4. Phospholipid hydroperoxides can react with metal ions, producing alkoxyl and peroxyl radicals. These radicals undergo intra-and inter-molecular reactions, producing aldehydes and truncated species, which can cause membrane rupture (47), and cell death by ferroptosis (48). Moreover, phospholipid hydroperoxides generate O_2_ (^1^Δ_g_) upon reacting with metal ions (30), heme proteins (32) and other oxidants such as hypochlorous acid (31). Notably, here we also demonstrate that plasmalogen hydroperoxyacetal reacts with metal ions (FeII and CeIV) producing O_2_ (^1^Δ_g_) through the Russell mechanism. This mechanism is supported by the detection of the corresponding alcohol and ketone products (Fig. 1B). In the case of pPE (p18:0-OOH/18:1), the conversion of hydroperoxyacetal to an alcohol yields a hemiacetal that cleaves, generating LPE and alpha-beta unsaturated fatty aldehyde. Interestingly, the conversion of hydroperoxyacetal to a ketone generates an ester-bonded acyl chain containing an alpha-beta unsaturated carbonyl group. The reactivity and biological consequences of this electrophilic phospholipid specie deserve further investigation.

Collectively, the results presented herein demonstrate that plasmalogen oxidation generates reactive intermediates with potential pro-oxidant and deleterious roles. These intermediates include triplet carbonyls, O_2_ (^1^Δ_g_), alpha-beta-unsaturated phospholipids and fatty aldehydes. Notably, excited molecules can propagate oxidative reaction in the absence of light, a phenomenon known as “dark photochemistry” (26, 41). Additionally, triplet carbonyls can cause deleterious effects in cells by transferring energy to other biomolecules, a process recently described for excited melanin, neurotransmitters, and hormones, leading to the formation of carcinogenic DNA modifications (49, 50). Hence, triplet carbonyl emerges as a product of plasmalogen oxidation with deleterious action. Furthermore, alpha-beta unsaturated lipids have the capacity to modify peptides, proteins, and other biomolecules. These aldehydes are susceptible to Michael addition reactions in the presence of nucleophiles, such as the sulfhydryl group of cysteines (51). For instance, 2-hexadecenal, an alpha-beta unsaturated long-chain aldehyde produced in sphingolipid enzymatic and non-enzymatic metabolism (52) has been reported to induce cytoskeletal reorganization and apoptosis in various cell types (53-55).

While typically regarded as antioxidants, our data shows that plasmalogens act as pro-oxidants when the vinyl-ether double bond reacts with singlet oxygen. This finding holds particular significance for understanding the biological roles of O_2_ (^1^Δ_g_) and stress responses in tissues and microorganisms exposed to sunlight (56), extending beyond the direct effects observed from UV radiation. This includes the dark reactions occurring post-light exposure in the skin, leading to DNA photoproducts long after UV exposure (26, 49) and the effects of other wavelength ranges, such as visible light acting on endogenous sensitizers (57). Overall, our study highlights potential pro-oxidant pathways of plasmalogens, revealing their involvement in excited species generation and reactive lipid production, thus contributing to a deeper understanding of oxidative processes in biological systems.

### Materials and Methods Plasmalogen photooxidation

pPE(p18/18:1) was photooxidized at a concentration of 1 mM in the presence of 0.3 mM methylene blue (MB) in CHCl_3_ (or CDCl_3_ for luminescence emission). PE (14:0/14:0) 0.1 mM was used as an internal standard. Samples were photooxidized with a LED lamp (632 nm, 41 ± 1 Wm-2) for 30 minutes in a -40°C bath (dry ice and ACN) while bubbling oxygen. Aliquots were collected 10, 20 and 30 min, and then diluted 100 times before LC-QTOF-MS/MS analysis. For comparison, a pPE mixture purified from bovine brain (app. 5 mM) was photooxidized in the presence of MB in CHCl_3_under the same conditions.

### Lipidomic analysis by LC-QTOF-MS/MS

Plasmalogen oxidation was comprehensively analyzed using lipidomic analysis conducted on an ESI-QTOF mass spectrometer (Triple TOF® 6600, Sciex, Concord, US) coupled with ultra-high performance liquid chromatography (UHPLC Nexera, Shimadzu, Kyoto, Japan). The samples were loaded into a CORTECS® (UPLC® C18 column, 1.6 μm, 2.1 mm i.d. × 100 mm) with a flow rate of 0.2 mL min^−1^ and the oven temperature maintained at 35 °C. For reverse-phase LC, mobile phase A consisted of water:acetonitrile (60:40, v/v), while mobile phase B composed of isopropanol:acetonitrile:water (88:10:2, v/v/v). Mobile phases A and B contained formic acid at a final concentration of 10 mM. Lipids were separated by a 20 min linear gradient as follows: from 40 to 100% B over the first 10 min., hold at 100% B from 10–12 min., decreased from 100 to 40% B during 12–13 min., and hold at 40% B from 13–20 min. Mass spectrometer was operated in the negative ionization mode with ion spray voltage of −4.5 and the cone voltage at −80 V. Additional parameters included: curtain gas set at 25 psi, nebulizer and heater gases at 45 psi and interface heater of 450 °C. MS1 and MS2 was acquired by Information Dependent Acquisition (IDA®) using the following parameters: m/z scan range 200–2000 Da; cycle time period of 1.05 s with 100 ms acquisition time for MS1 scan and 25 ms acquisition time to obtain the top 36 precursor ions. The LC-MS/MS data was acquired by Analyst ® 1.7.1 and data was analyzed with PeakView®. Lipid molecular species were manually identified based on their exact masses, specific fragments and/or neutral losses. A maximum error of 5 mDa was defined for the attribution of the precursor ion. After identification, the area of lipid species was obtained by MS1 data using MultiQuant®. Each peak integration was carefully inspected for correct peak detection and accurate area determination. PE(14:0/14:0) was used as an internal standard for pPE data normalization.

### Quantification of plasmalogen hydroperoxides

For the quantification of plasmalogen hydroperoxides the method based on triphenylphosphine (TPP) reaction with hydroperoxides was employed(58). Briefly, 25 μL of the photooxidized pPE solution was added to a 100 μL insert. The solvent was removed using a nitrogen stream, and 50 μL of 5 mM TPP in ACN was added. The mixture was incubated for 30 minutes in the dark at room temperature under stirring at 300 rpm. Subsequently, TPP and its oxide (TPPO) were analyzed by ultra-high performance liquid chromatography (UHPLC Nexera, Shimadzu, Kyoto, Japan) using a CORTECS® (UPLC® C8 column, 1.6 μm, 2.1 mm i.d. × 100 mm). The flow rate was set at 0.3 mL/min and the oven temperature maintained at 40 °C. The injection volume was 5 μL, and the UV detection wavelength was 220 nm. UHPLC separation was done with acetonitrile (solvent B) and water (solvent A) using the following steps: a linear gradient from 40% to 100% B over 0-1 min, holding at 100% B from 1–2 min, decreasing from 100% to 40% B during 2–2.1 min, and holding at 40% B from 2.1–4.5 min. A calibration curve was constructed with tert-butylhydroperoxide or LAOOH.

### Plasmalogen hydroperoxide incubation with metal ions

Incubations of plasmalogen hydroperoxides with metal ions were done by mixing 5 μM pPE(p18-OOH/18:1) or PE (18:1/18:1)-OOH with CeIV or FeII in CHCl_3_:methanol:water (90:9:1 v/v/v) at 37°C for 1 hour. Reaction products were analyzed by LC-QTOF-MS/MS using PE (14:0/14:0) at 0.5 μM as internal standard. The LC system setup included a valve that was used to discard the initial 5 minutes to avoid interference from metal ions (CeIV or FeII) in the mass spectrometry data acquisition. For singlet oxygen light emission measurement in the NIR region, we added 308 μL of pPE(p18-OOH/18:1) or LA-OOH in CDCl_3_ in a cuvette. Then 92 μL of a MeOD:D_2_O (914:6; v/v) solution containing metal ions (CeIV or FeII) was infused into the solution within 60 seconds under stirring, resulting in a final solution with 5 μM hydroperoxide and 75 μM metal ion (CeIV or FeII).

### Fatty aldehyde derivatization

For the analysis of alpha-beta unsaturated lipids, 100 μM of photooxidized pPE with or without FeII/CeIV was incubated with 5 mM CSH in ACN with 0.1% TEA for 2 hours at room temperature before incubation with CHH. Derivatized fatty aldehyde samples were analyzed by ultra-high performance liquid chromatography (UHPLC Nexera, Shimadzu, Kyoto, Japan). Aliquots of 5μL of the sample was injected into a reversed-phase column water cortex C8 (100 × 4.6 mm, 1.7 μm particle size) with a flow rate of 0.2 mL min^−1^ and the oven temperature maintained at 35 °C and RF-10Axl fluorescence detector. The HPLC mobile phase consisted of water with 0.1% formic acid (A) and methanol with 0.1% formic acid (B), and the flow rate was 0.3 mL/min. The separation of the fluorescent adducts was done using the following condition: 60% B for 2 min, 60-100% in 9 min, 92% B for 3 min, and 100-60% B in 1 min and 60% for 3min. The excitation and emission wavelengths were fixed at 450 and 468 nm.

### Fatty aldehyde analysis by LC-QTOF-MS/MS

The aldehydes derivatized with CHH were analyzed by LC-QTOF (Triple TOF® 6600, Sciex, Concord, US) interfaced with an ultra-high performance liquid chromatography (UHPLC Nexera, Shimadzu, Kyoto, Japan). The chromatographic conditions were the same as for the fluorescence analysis. To avoid ion suppression due to excess CHH probe, we used a valve to discard the first initial 5 min (**Fig. 6S E**) in the LC-MS analysis. All aldehydes entering the MS were analyzed in positive mode (**Fig. 6S F**). The final derivation conditions were: 50 μM aldehydes or 50 μM pPE, 1 mM CHH in isopropanol with 0.1% formic acid.

### Excited triplet carbonyl light emission measurements in the visible region

All reactions were carried out in a quartz tube under constant stirring at room temperature (final volume = 600 μL). The light emission in the visible region was immediately recorded by a FLSP 920 photon counter (Edinburgh Instruments, Edinburgh, UK) with a PMT Hamamatsu detector R9110, maintained at −20 °C by a PMT cooler CO1 (Edinburgh Instruments). 1 mM of pPE(p18/18:1) or PE (18:1/18:1) photo-oxidized at -40°C in CDCl_3_ was added to a cuvette and naturally warmed to room temperature. For carbonyl enhancer experiments, DBA was added before the reading at a final concentration of 10 mM. For chemical trapping, DPA was added prior to the reading at a final concentration of 10 mM and incubated for 1 hr at room temperature, then analyzed by HPLC at 210 nm(30).

### Singlet molecular oxygen light emission measurements in the near-infrared region

The monomolecular light emission of O_2_ (^1^Δ_g_) was measured using two photocounting apparatus developed in our laboratory, equipped with a monochromator capable of selecting emissions in the near-infrared (NIR) region (950-1400 nm).The NIR spectra were recorded by a FLSP 920 photon counter (Edinburgh Instruments, Edinburgh, UK) consisting of a H10330A-45 NIR-PMT detector (Hamamatsu city, Japan) that is coupled to a thermoelectric cooled module apparatus maintained at -60 °C to reduce the dark current. The power was provided by a high voltage DC power supply, and the applied potential was set to –0.8 kV. The light emitted from the sample was processed through a monochromator (TMS/DTMS300, Edinburgh Analytical Instruments, UK) equipped with a diffraction grating capable of selecting wavelengths in the infrared region. The phototube output was connected to the computer, and the signal was acquired. The monochromator was controlled, and the data was acquired using the F-900 version 6.22 software (Edinburgh Analytical Instruments, Livingston, UK). The NIR light emissions kinetics at 1270 nm was done in a second photocounting apparatus using a band pass filter placed between the cuvette and the photomultiplier. All sample components were mixed and poured into a glass cuvette (35 × 7 × 55 mm) maintained at 25 °C.

## Data Availability

Raw mass spectrometry analysis files (wiff data) will be available at EMBL-EBI MetaboLights database (DOI: 10.1093/nar/gkad1045, PMID:37971328) with the identifier MTBLS9564.

Any additional data required to reanalyze the data will be available from the corresponding author upon request.

## Acknowledgments

We thank Adriano Britto Chaves-Filho and Marcos Y. Yoshinaga for their support with mass spectrometry and lipidomic analysis. We thank Miyamoto Lab members and Cepid-Redoxoma group for their constant support and discussions. We thank Erick L. Bastos and Etelvino Bechara for helpful discussions. This study was supported by Fundação de Amparo à Pesquisa do Estado de São Paulo [FAPESP, CEPID-Redoxoma, 13/07937-8], Conselho Nacional de Desenvolvimento Científico e Tecnológico [CNPq 313926/2021-2 to S.M., 302120/2018-1 to P.D.M.], Coordenação de Aperfeiçoamento de Pessoal de Nível Superior [CAPES, Finance Code 001] and Pro-Reitoria de Pesquisa da Universidade de São Paulo [PRPUSP]. The PhD scholarship of R.L.F was supported by FAPESP [17/16140-7].

## Author Contributions

R.L.F. and S.M. designed research; R.L.F., F.M.P., H.C.J., K.C.F., L.R.D. performed research; R.L.F., F.M.P., M.S.B, P.D.M., and S.M., analyzed data; and R.L.F, M.S.B, P.D.M., and S.M. wrote the paper.

## Supporting information available

Supporting text (Materials)

Figures S1 to S7

Tables S1

SI References

