## Supporting information for "Plasmalogen oxidation induces the generation of excited molecules and electrophilic lipid species"

**This PDF file includes:**

Supporting text (Materials)
Figures S1 to S7
Tables S1
SI References

**Supporting Information Text**

**Materials**

**From Sigma Aldrich (Saint Louis, MO, USA):**

- 7-(Diethylamino)coumarin-3-carbohydrazide (CHH, cod: 36798, Lot: # BCCF3603)
- trans,trans-2,4-Nonadienal (cod: 61410, Lot: # BCBR3933V)
- trans,trans-2,4-Decadienal (cod: 90628, Lot: # BCBW7178)
- Tridecanal (cod: 269239, Lot: # MKCB9020)
- Hexenal (cod: 76717, Lot: #BCBW3602)
- 9,10-Dibromoanthracene (cod: D38855, Lot: # 02503EH)
- 9,10-Diphenylanthracene (cod: D205001)
- Methanol-d4 (MeOD, cod: 441384, Lot: 01006TH)
- Ammonium cerium(IV) nitrate (CeIV, cod: 215473)
- Iron(II) sulfate hydrate (FeII, cod: 307718, Lot:STBJ9273)
- Deuterium oxide (D2O, cod: 151882, Lot: MKBP9326V)
- 7-Mercapto-4-methylcoumarin (CSH, cod: 63759, Lot: BCCD3153)
- Tert-Butyl hydroperoxide solution (cod: 458139, Lot: 00112MH)

**From Avanti Polar Lipids (Alabaster, AL, USA):**

- 1-(1Z-octadecenyl)-2-oleoyl-sn-glycero-3-phosphoethanolamine (pPE(p18/18:1), cod: 852758P, Lot: 852758P -10MG-A-024)
- 1,2-dimyristoyl-sn-glycero-3-phosphoethanolamine (PE(14:0/14:0), cod: 850745P, Lot: 5738P-NA-070)
- (2E)-hexadecenal or 16:1 aldehyde (Ald 16:1 Δ2, cod: 857459P, Lot: 857459P -1MG-D-010)
- 1-palmitoyl-2-(9'-oxo-nonanoyl)-sn-glycero-3-phosphocholine (ALDOPC, cod: 870605P, Lot: 870605P-1MG-F-014)

**From Cayman Chemicals (Ann Arbor, MI):**

- 4-hydroxy-hexenal (HHE, Cod: 32060, Lot: 0411894-9)
- 4-hydroxy-nonenal (HNE, Cod: 32100, Lot: 0411894-9)
- Hexadecanal (Ald 16:0, cod: 9001996, Lot: 050947-4)

**From Tokyo Chemical Industry (Nihonbashi-honcho, Chuo-ku, Tokyo, Japan):**

- Heptadecanal (Ald (17:0), cod: H1295, Lot: MBUJA-MQ)
- Pentadecanal (Ald (15:0), cod: P1869, Lot: KODWM-IE)

1,4-Dimethylnaphthalene (DMN) endoperoxide (DMNO2) was prepared by UVA irradiation of DMN/methylene blue and then quantified spectrophotometrically (1).

Linoleic acid hydroperoxide (LA-OOH) was synthesized following the method described by Miyamoto et al (2).

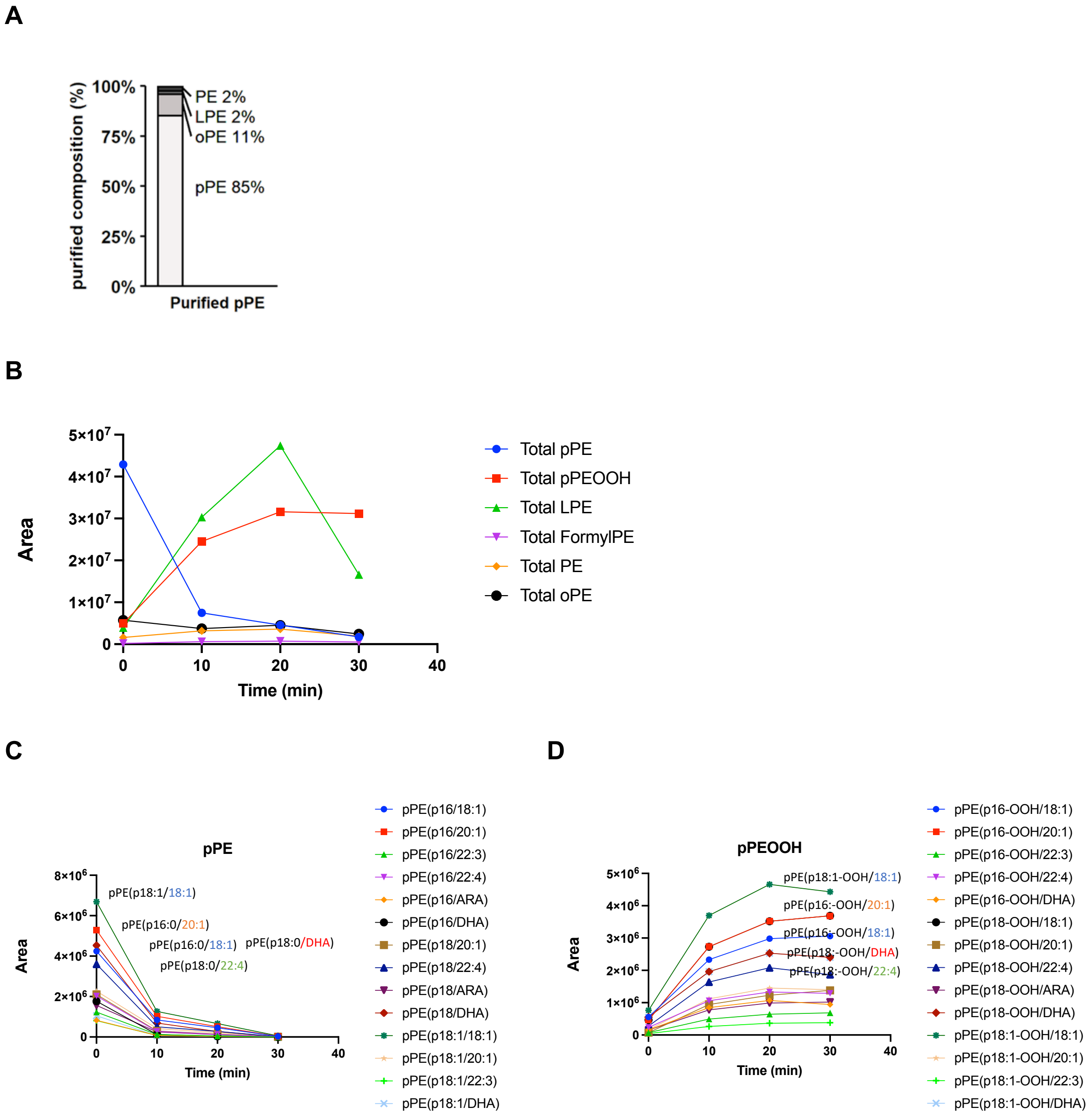


**Fig. S1.** (A) Lipidomic analysis showing the composition of PE plasmalogen purified from bovine brain.

(B) Total amounts of plasmalogen species consumed by photooxidation in the presence of methylene (blue) and the generation of hydroperoxyacetal intermediates (red), LPE (green), Formyl-PE (violet).

(C) Consumption of pPE species by photooxidation

(D) Generation of pPE hydroperoxyacetal intermediates

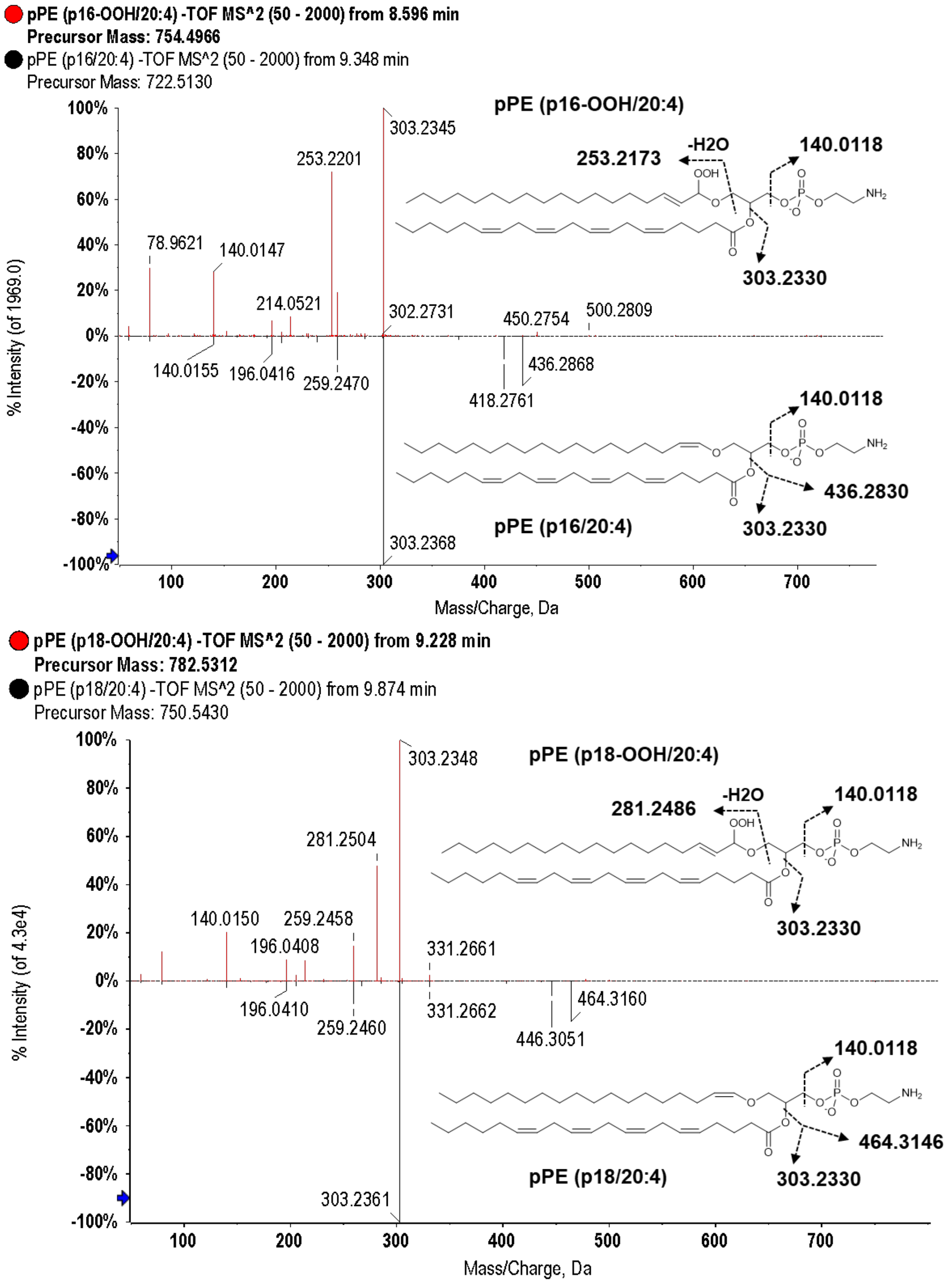


**Fig. S2**. Comparative MS/MS spectra of non-oxidized plasmalogen and its corresponding sn-1 ether hydroperoxides: (Top) pPE(p16-OOH/20:4) and pPE(p16/20:4); (Lower) pPE(p18-OOH/20:4) and pPE(p18/20:4). These plasmalogen species were detected during brain plasmalogen mixture photooxidation in the presence of methylene blue.

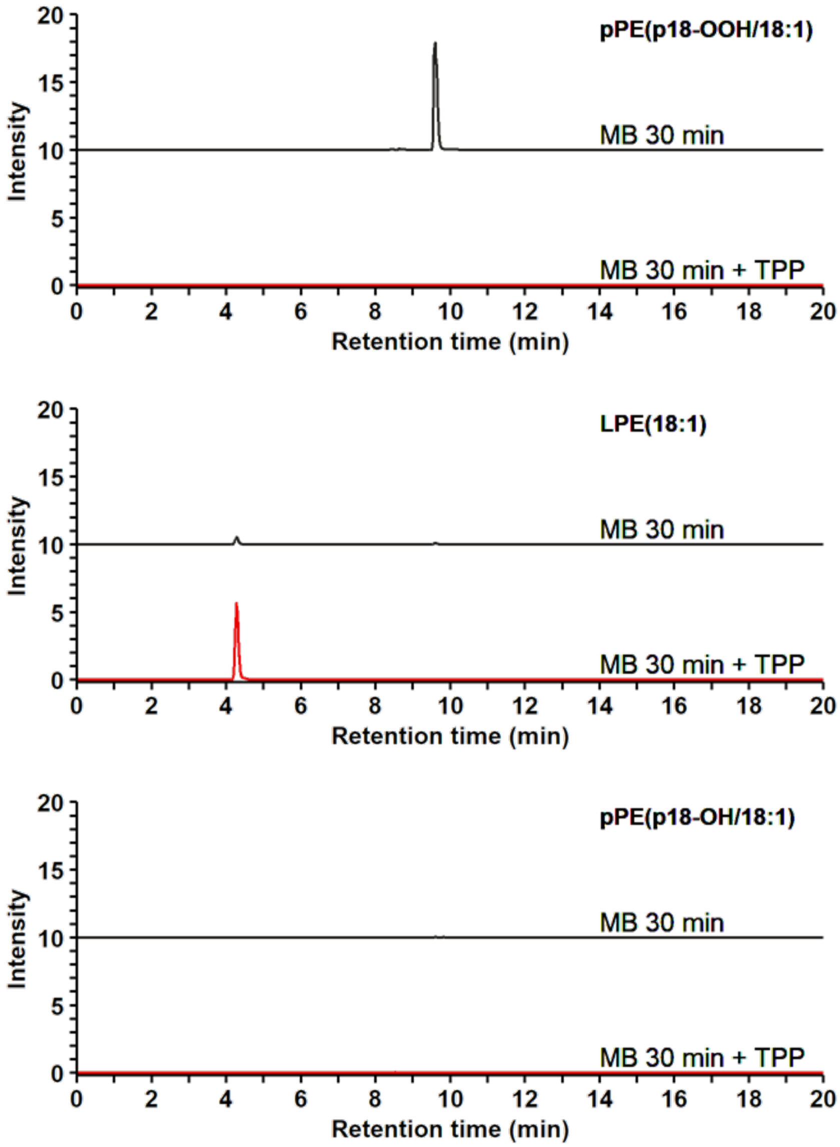


**Fig. S3.** Extracted ion chromatograms (XIC) of pPE(p18/18:1) photooxidised in the presence of MB for 30 minutes, with (red line) or without (black line) TPP: (Top) XIC m/z:760.55-760.56 from pPE (p18-OOH/18:1); (Middle) XIC m/z:478.29-478.230 from LPE (18:1); (Bottom) XIC m/z: 744.55-744.56 from pPE (p18-OH/18:1).

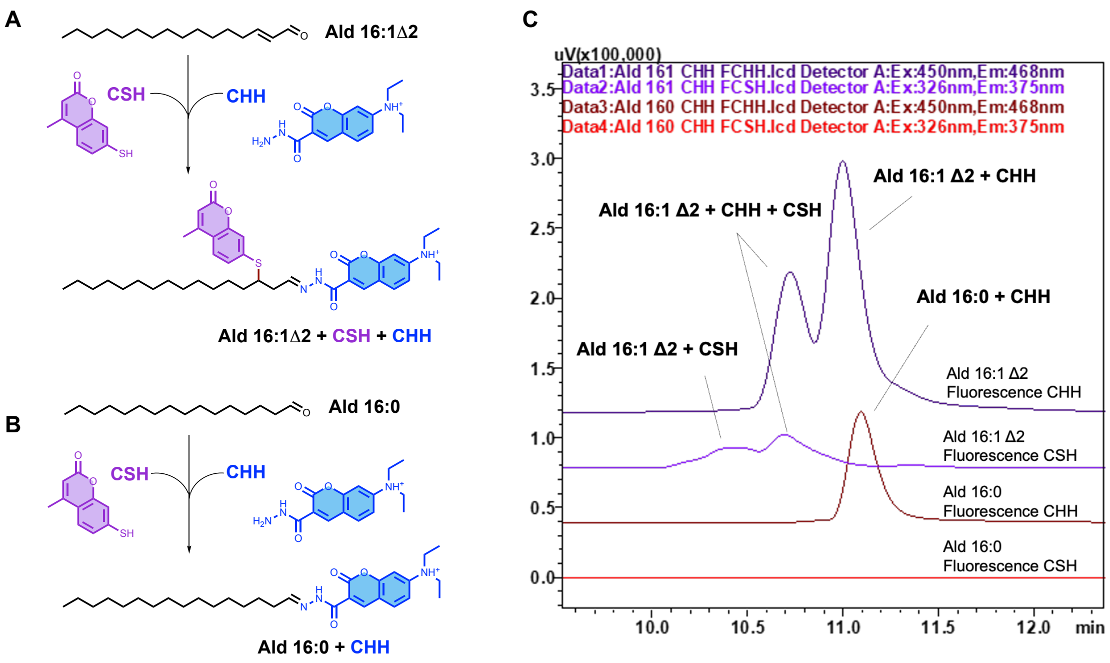


**Fig. S4. Fatty aldehyde derivatization with CSH and CHH probes**. Derivatization of (A) Ald (16:1 Δ2); and (B) Ald (16:0) with CSH and CHH probes. (C) HPLC chromatograms of Ald(16:1Δ2)-CHH-CSH and Ald(16:0)-CHH adducts detected by fluorescence analysis of CSH (excitation: 326 nm and emission: 375 nm) and CHH (excitation: 450 nm and emission: 468 nm).

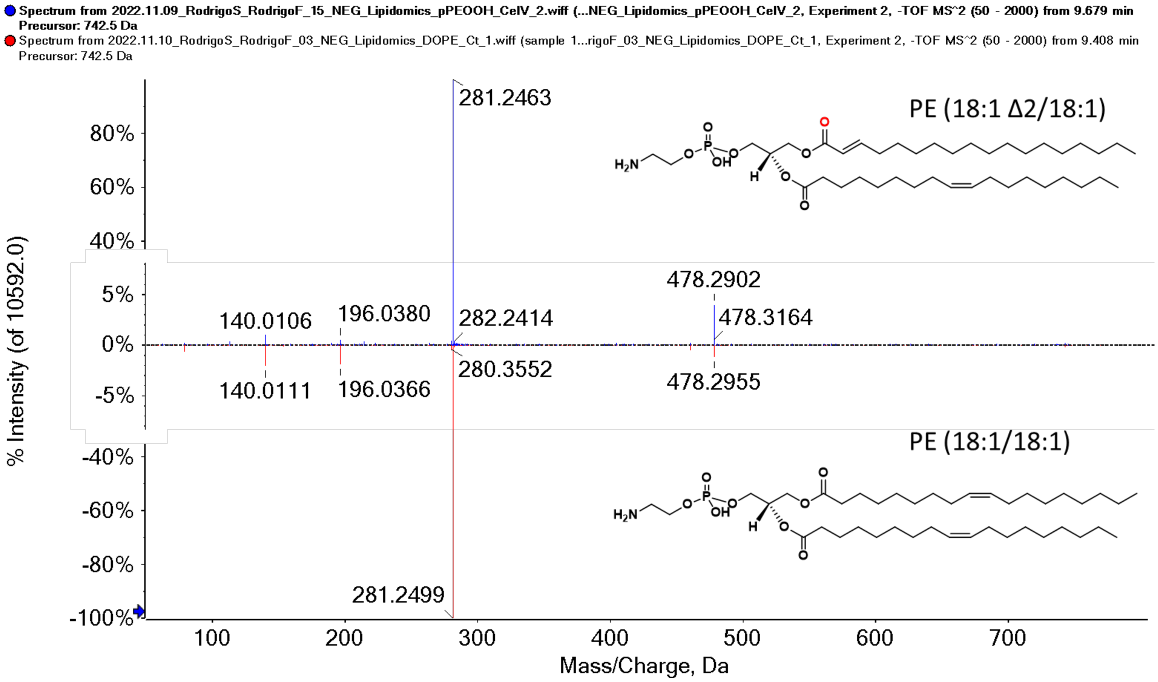


**Fig. S5.** Comparative MS/MS spectra of commercial PE (18:1/18:1) and PE (18:1Δ2/18:1) produced by pPE (p18/18:1) photooxidation.

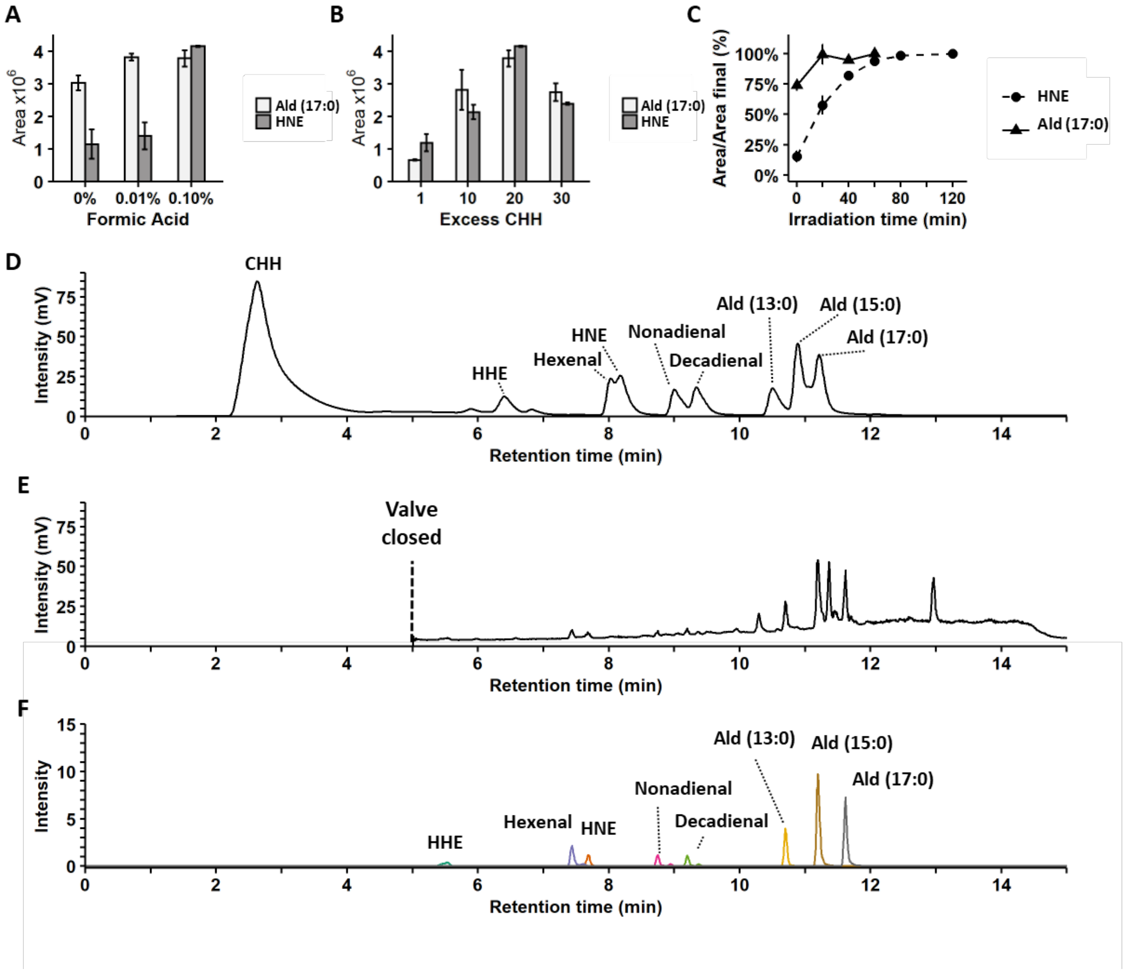


**Fig. S6. Optimization of aldehyde derivatization conditions using the CHH probe.**
(A) Effect of formic acid in on Ald(17:0) and HNE aldehyde derivatization. (B) Effect of CHH concentration in relation to Ald(17:0) aldehydes or HNE derivatization in the presence of 0.1% formic acid. (C) Time-course analysis of Ald(17:0) or HNE derivatization with 20x molar excess of CHH and 0.1% formic acid. (D) Chromatogram a aldehyde standard mix incubated with 20x molar excess of CHH and 0.1% formic acid. (E) Total ion chromatogram of aldehydes derivatized with CHH. (F) Extracted ion chromatograms of aldehydes derivatized with CHH. The chromatography method was optimized so that short-chain aldehydes [4-hydroxyhexenal, HHE; 4-hydroxynonenal, HNE; Hexenal, Nonadienal and Decadienal] were separated from long-chain fatty aldehydes [Ald (13:0), Ald (15:0), Ald (17:0)].

Fatty aldehyde analysis method was adapted from Mansano et al.(3) using the 7-(Diethylamino)coumarin-3-carbohydrazide (CHH). We determined the optimal concentration of formic acid to catalyze the derivatization reaction as 0.1% v/v (**Fig S6A**). Additionally, we established that the optimal aldehyde derivatization is achieved with a 20-fold molar excess of the probe relative to aldehydes (**Fig S6B**) and the ideal reaction time of 120 min (**Fig S6C**).

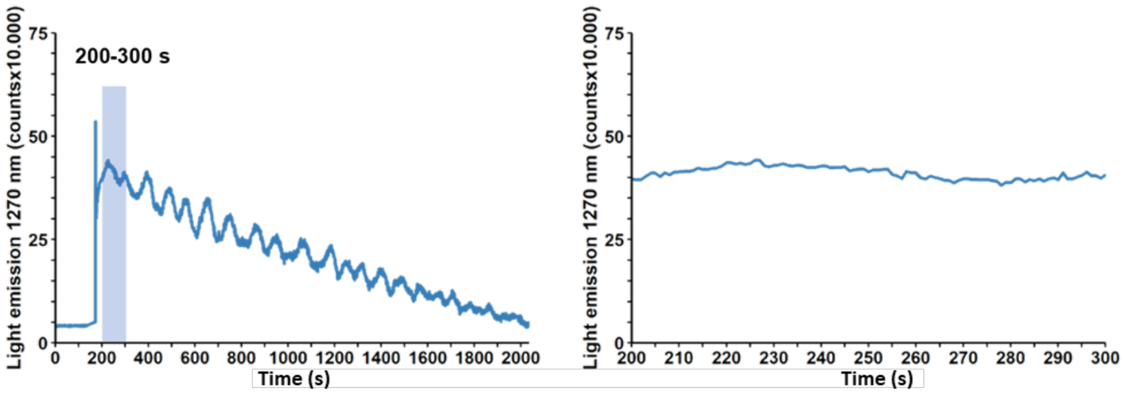


**Fig. S7.** NIR light emission at 1270 nm obtained for 116 μM DMNO2 decomposition in 400 μL $CDCl_3$:MeOD(d5):$D_2O$ solution 308:91.4:0.6 v/v/v (A) Light emission kinetics recorded until all DMNO2 was decomposed (B) Light emission recorded from 200 to 300s was used for the estimation of singlet oxygen yields.

**Table S1.** Relative percentage of pPE molecular species detected before and after 30 min photooxidation with methylene blue at -40C. The time dependent consumption of total pPE is shown in Figure S1. Samples were analyzed by LC-Q-TOF-MS/MS and the peak areas of all detected pPE species were normalized against PE (14:0/14:0) internal standard. Normalized area ratio of all pPE species were summed and presented as relative %. The analysis encompasses the time points, namely the initial condition (0 min), the final point after 30 minutes of photooxidation (30 min) without and with the addition of triphenylphosphine to reduce the hydroperoxides (30 min + TPP). Data are expressed as percentages relative to the initial condition (0 min).

| Component Name | 0 min | 30 min | 30 min + TPP |
|---|---|---|---|
| pPE(p16/18:1) | 7.41 | 0.06 | 0.03 |
| pPE(p16/20:1) | 9.23 | 0.07 | 0.03 |
| pPE(p16/20:3) | 0.30 | 0.00 | 0.00 |
| pPE(p16/20:4) | 1.44 | 0.00 | 0.00 |
| pPE(p16/22:3) | 2.15 | 0.00 | 0.00 |
| pPE(p16/22:4) | 3.54 | 0.02 | 0.01 |
| pPE(p16/22:6) | 3.09 | 0.00 | 0.00 |
| pPE(p17/18:1) | 1.14 | 0.00 | 0.00 |
| pPE(p17/20:1) | 0.57 | 0.00 | 0.00 |
| pPE(p17/22:4) | 0.48 | 0.00 | 0.00 |
| pPE(p18/16:0) | 0.12 | 0.01 | 0.00 |
| pPE(p18/18:1) | 0.91 | 0.22 | 0.16 |
| pPE(p18/20:1) | 3.67 | 0.02 | 0.00 |
| pPE(p18/20:4) | 2.73 | 0.01 | 0.01 |
| pPE(p18/22:1) | 0.16 | 0.10 | 0.03 |
| pPE(p18/22:4) | 6.28 | 0.02 | 0.00 |
| pPE(p18/22:5) | 0.34 | 0.00 | 0.00 |
| pPE(p18/22:6) | 7.91 | 0.02 | 0.01 |
| pPE(p18/24:3) | 0.16 | 0.00 | 0.00 |
| pPE(p18:1/18:1) | 11.67 | 0.09 | 0.05 |
| pPE(p18:1/19:1) | 0.42 | 0.00 | 0.00 |
| pPE(p18:1/20:1) | 3.97 | 0.05 | 0.03 |
| pPE(p18:1/20:2) | 0.77 | 0.07 | 0.06 |
| pPE(p18:1/20:3) | 0.22 | 0.00 | 0.01 |
| pPE(p18:1/21:1) | 0.32 | 0.00 | 0.00 |
| pPE(p18:1/22:1) | 0.81 | 0.15 | 0.10 |
| pPE(p18:1/22:3) | 1.42 | 0.03 | 0.01 |
| pPE(p18:1/22:6) | 1.84 | 0.00 | 0.00 |
| pPE(p18:1/24:1) | 0.06 | 0.00 | 0.00 |
| pPE(p18:1/24:3) | 0.36 | 0.01 | 0.00 |
| pPE(p18:1/24:4) | 0.13 | 0.00 | 0.00 |
| pPE(p16-OOH/18:1) | 0.97 | 5.34 | 0.00 |

| | | | |
|---|---|---|---|
| pPE(p16-OOH/20:1) | 0.81 | 6.43 | 0.00 |
| pPE(p16-OOH/22:3) | 0.12 | 1.19 | 0.00 |
| pPE(p16-OOH/22:4) | 0.41 | 2.25 | 0.00 |
| pPE(p16-OOH/22:6) | 0.27 | 1.65 | 0.00 |
| pPE(p17-OOH/18:1) | 0.42 | 0.08 | 0.00 |
| pPE(p18:1-OOH/18:1) | 1.34 | 7.73 | 0.00 |
| pPE(p18:1-OOH/20:1) | 0.28 | 2.45 | 0.00 |
| pPE(p18:1-OOH/20:2) | 0.03 | 0.31 | 0.00 |
| pPE(p18:1-OOH/20:4) | 0.13 | 1.96 | 0.00 |
| pPE(p18:1-OOH/21:1) | 0.01 | 0.16 | 0.00 |
| pPE(p18:1-OOH/22:1) | 0.03 | 0.42 | 0.00 |
| pPE(p18:1-OOH/22:2) | 0.01 | 0.17 | 0.00 |
| pPE(p18:1-OOH/22:3) | 0.08 | 0.67 | 0.00 |
| pPE(p18:1-OOH/22:6) | 0.97 | 4.19 | 0.00 |
| pPE(p18-OOH/18:1) | 0.81 | 6.43 | 0.00 |
| pPE(p18-OOH/20:1) | 0.13 | 2.42 | 0.00 |
| pPE(p18-OOH/20:4) | 0.27 | 1.79 | 0.00 |
| pPE(p18-OOH/22:1) | 0.01 | 0.13 | 0.00 |
| pPE(p18-OOH/22:3) | 0.06 | 1.02 | 0.00 |
| pPE(p18-OOH/22:4) | 0.49 | 3.27 | 0.00 |
| pPE(p18-OOH/22:5) | 0.02 | 0.15 | 0.00 |
| pPE(p18-OOH/22:6) | 0.97 | 4.19 | 0.00 |
| pPE(p18:1-OOH/18:1-OOH) | 0.00 | 1.47 | 0.00 |
| pPE(p18:1-OOH/22:4-OOH) | 0.00 | 0.16 | 0.00 |
| pPE(p18-OOH/22:4-OOH) | 0.00 | 0.22 | 0.00 |
| LPE(16:0) | 0.09 | 0.55 | 1.25 |
| LPE(16:1) | 0.02 | 0.06 | 0.15 |
| LPE(18:0) | 0.13 | 0.34 | 0.42 |
| LPE(18:1) | 3.65 | 13.22 | 24.39 |
| LPE(20:1) | 0.99 | 4.60 | 9.48 |
| LPE(20:3) | 0.11 | 0.53 | 1.34 |
| LPE(20:4) | 0.33 | 3.07 | 4.97 |
| LPE(22:1) | 0.00 | 0.00 | 0.00 |
| LPE(22:2) | 0.04 | 0.17 | 0.44 |
| LPE(22:3) | 0.04 | 0.15 | 0.42 |
| LPE(22:4) | 1.27 | 3.82 | 7.77 |
| LPE(22:5) | 0.09 | 0.29 | 1.30 |
| LPE(22:6) | 0.11 | 2.23 | 8.11 |
| PE(Formyl/18:1) | 0.20 | 0.64 | 0.52 |
| PE(Formyl/20:1) | 0.00 | 0.01 | 0.01 |
| PE(Formyl/20:4) | 0.04 | 0.10 | 0.09 |

| | | | |
|---|---|---|---|
| PE(Formyl/22:4) | 0.06 | 0.16 | 0.12 |
| PE(Formyl/22:5) | 0.01 | 0.03 | 0.10 |
| oPE(o16/20:1) | 1.44 | 1.43 | 1.18 |
| oPE(o16/22:3) | 0.12 | 0.13 | 0.11 |
| oPE(o18/16:1) | 0.72 | 0.62 | 0.51 |
| oPE(o18/20:1) | 0.92 | 0.80 | 0.59 |
| oPE(o18/22:1) | 0.08 | 0.07 | 0.06 |
| oPE(o18/22:4) | 0.84 | 0.60 | 0.47 |
| oPE(o18/22:6) | 3.72 | 0.50 | 0.39 |
| PE(16:0/18:1) | 0.24 | 0.23 | 0.28 |
| PE(18:0/18:1) | 0.46 | 0.68 | 0.59 |
| PE(18:0/20:1) | 0.06 | 0.08 | 0.06 |
| PE(18:0/20:4) | 0.37 | 0.29 | 0.39 |
| PE(18:0/22:4) | 0.12 | 0.20 | 0.15 |
| PE(18:0/22:5) | 0.02 | 0.10 | 0.08 |
| PE(18:0/22:6) | 1.02 | 0.79 | 0.79 |
| PE(18:1/18:1) | 0.32 | 0.33 | 0.29 |
| PE(18:1/18:2) | 0.08 | 0.31 | 0.25 |
| PE(18:1/20:1) | 0.01 | 0.06 | 0.04 |
